# BNIP3-mediated mitophagy boosts the competitive dominant growth of lenvatinib resistant cells via reprogramming energy metabolism in HCC

**DOI:** 10.1101/2023.07.12.548688

**Authors:** Sikai Wang, Hongxia Cheng, Miaomiao Li, Haoran Wu, Shanshan Zhang, Dongmei Gao, Yilan Huang, Kun Guo

**Author notes:** Correspondence (K.G.). These authors contributed equally (Sikai Wang, Hongxia Cheng, Miaomiao Li).

## Abstract

Although increasing studies has demonstrated that cell competition widely involved in the growth and homeostasis of multicellular organisms is closely linked to tumorigenesis and development, the mechanistic contributions to the association between tumor cell competition-driven heterogeneity and drug resistance remains ill-defined. In our study, lenvitinib-resistant hepatocellular carcinoma (HCC) cells display obviously competitive growth dominance against sensitive cells through reprogramming energy metabolism. Mechanistically, when BCL2 interacting protein3 (BNIP3) overexpression activates mitophagy activity in lenvatinib-resistant HCC cells, energy imbalance signal caused by reduced mitochondrial oxidative phosphorylation levels provokes the phosphorylation of AMP-activated protein kinase (AMPK) sensor; subsequently, enabled AMPK specifically targets enolase 2 (ENO2) to enhance glycolysis and eventually promots the competitive capacity and dominant growth. Of note, BNIP3 deficiency shows certain inhibition of cell competition outcome. Our findings emphasize a vital role for BNIP3-AMPK-ENO2 signaling in maintaining the competitive outcome of lenvitinib-resistant HCC cells via regulating energy metabolism; meanwhile this work recognaizes BNIP3 as a promising target to overcome HCC drug resistance.

## Introduction

Cell competition, a vital selection and quality control mechanism for maintaining tissue or organ development and homeostasis in multicellular organisms, executes the process of detecting and excluding cells regarded as less fit (loser) by the fitter compeer (winner), characterized by the survival of the fittest (van Neerven & Vermeulen, 2023). During embryogenesis, depending on the differential expression of certain genes, (for example, GATA6, YAP and MYC), the optimal and fittest pluripotent cells of epiblast are outcompeted and selected for differentiating into various cell types and eventually form the complete organism via apoptosis pathway (Chazaud & Yamanaka, 2016; Claveria *et al*, 2013; van Neerven & Vermeulen, 2023). Of note, metabolic fitness can function as the trigger of active cell competition. A study on mouse embryo has showed that active cell competition eliminated epiblast cells with mitochondrial dysfunction (Lima *et al*, 2021). As the growth of the organism, cell competition is primarily involved in tissue or organ development and homeostasis. For example, to maintain the integrity of skin, the less fit epidermal stem cells with low expression of COL17A1 are pushed out mechanically and replaced by the high expression population (Liu *et al*, 2019). In this period, importantly, cells originated from different tissues or organs are subjected to mutate constantly and, those with gene mutation generally representing deregulated signaling pathways such as p53 signaling and WNT signaling in cancer can both outcompete as well as lose during cell competition. These phenomena suggest that cell competition is engaged in tumorigenesis (Baker, 2020; Parker *et al*, 2021; Vishwakarma & Piddini, 2020). Based on epithelial defence against cancer (EDAC), unmutated epithelial cells mechanically eliminate aberrant cells accompanied by Tp53 mutation, Kras mutation or Src mutation through apical extrusion (Kajita *et al*, 2010; Prior *et al*, 2020; Watanabe *et al*, 2018). Gradually increasing evidences, however, suggest that potentially tumorigenic cells from different tissue checkmate their opponent and expanse rapidly via imitating and obtaining a fitness dominance, eventually promoting tumor initiation and development. DNMT3A mutation endows hematopoietic stem cells (HSCs) with fitness advantage and prompts them to defeat unmutated HSCs after transplantation, thereby acquiring a major population dominance (Shlush *et al*, 2014). Through forming a favorable microenvironment of crypt colonization, Apc, Kras, or PIK3CA mutated-intestinal stem cells (ISCs) serving as supercompetitors actively compete and remove normal ISCs, eventually driving oncogenesis (2021; Flanagan *et al*, 2021; van Neerven *et al*, 2021). Recently, multiple novel anti-tumor drugs and radiotherapy that target cell competition (for example, facilitating EDAC and inhibiting mutant clones) are developed and applied in clinical treatment (Fernandez-Antoran *et al*, 2019; Sato *et al*, 2020; van Neerven *et al*., 2021). Unfortunately, drug resistance based on spatial- and temporal-heterogeneity, the inevitable challenge, has remarkably impeded their clinical efficacy. To overcome drug resistance, most focuses still simply lies in targeting oncogenes or boosting antitumor immunity. However, tumor heterogeneity is analogous to a knotted Gordian Knot, in which genetic changes and evolutionary selection confer it with multifaceted phenotypes, suggesting that it remains difficult to overcome drug resistance by focusing on previously non-interactive strategies, although they indeed obtain certain favorable effects. As a result, a tumor heterogeneity-based treatment strategy for drug resistance reversal would be a potential approach. Nevertheless, the mechanism between cell competition and drug resistance in tumors remains so far uncleared.

Therefore, in the present study, we chose hepatocellular carcinoma (HCC), one of the malignant tumors with high indidence and poor prognosis in China, and demonstrated the mechanism by which cell competition drove the dominant growth of lenvatinib resistant HCC cells; moreover, BNIP3-mediated mitophagy played a vital role in outcompeting and eliminating sensitive cells by lenvatinib resistant cells in HCC. In addition, mitophagy was identified to facilitate shifting energy production from mitochondrial oxidative phosphorylation to glycolysis via BNIP3-AMPK-ENO2 signaling to maintain perpetually HCC lenvatinib-resistant cells’ competitive dominance. Altogether, our study uncovered a promising approach to overcome drug resistance of HCC basing on tumor heterogeneity.

## Results

### Cell competition drives the dominant growth of lenvatinib-resistant cells in HCC

To characterize the biological behaviour conferred by lenvatinib resistance of HCC cells, we established successfully the lenvatinib-resistant HCC cell strain (Huh7R; IC50 = 40.91μm) by culturing lenvatinib-sensitive cell (Huh7; IC50 = 3.45μm) with progressively increased doses of lenvatinib (3-30 μm) for six months (**Fig EV1A-D; Video 1-4**); moreover, we stably tagged Huh7 with the tracer mcherry (Huh7m), which displayed significantly no difference from Huh7 in terms of cell migration, proliferation, apoptosis, and intracellular reactive oxygen species (ROS) level (**Fig EV1E-M**). Thereafter, we constructed a HCC cell competition in vitro model by coculturing Huh7R and Huh7m at a 1:1 ratio (defined cell competition group [CC group], in which Huh7R and Huh7m were respectively termed as CCHuh7R and CCHuh7m); meanwhile, non-competition group [NCC group] was set up by coculturing Huh7 and Huh7m at a 1:1 ratio, in which Huh7 and Huh7m were respectively termed as NCCHuh7 and NCCHuh7m. In the models, cell competition phenotype was evaluated based on the combination of flow cytometric analysis and visualization with high content imaging (**Fig 1A**). In cell competition group, CCHuh7m suffered growth inhibition and cell death, while CCHuh7R obtained concomitantly increased proliferation during in vitro-coculture for 48 hours; no significant cell competition phenomena was observed in non-competition group (**Fig 1B; Video 5 and Video 9**). The outcompeting of lenvatinib-resistant HCC cells against sensitive cells was further verified in cell-derived xenograft tumors in nude mice (**Fig 1C**). We preliminarily speculate that these biological characteristics reflect a common conditional foundation. To prove that the elimination of CCHuh7m and the significant difference in the proliferation rate of equivalent cells indeed resulted from the occurrence of cell competition, we compared the growth rates of a variety of cells in CC group (CCHuh7R, CCHuh7m), NCC group (NCCHuh7, NCCHuh7m) and alone culture group (Huh7, Huh7m, Huh7R) (**Fig 1D; Video 6-10**). Although there existed a different baseline of growth rate that Huh7R grew faster than Huh7m or Huh7 (Gomes *et al*, 2022; Shen *et al*, 2013), as expected, CCHuh7R proliferated significantly faster than Huh7R and NCCHuh7, while CCHuh7m grew slower than Huh7m or NCCHuh7m, even experienced mortality (**Fig 1E**). However, there was no proliferation difference among NCCHuh7m, NCCHuh7, Huh7m and Huh7. Moreover, the population ratios (m+/m-) displayed a remarkably descending trend in NCC group, single Huh7m mixed with single Huh7R group and CC group at indicated 24h/48h, or specifically in CC group over a time course of coculture, further validating the above results (**Fig 1F and G**).

**Figure 1.**
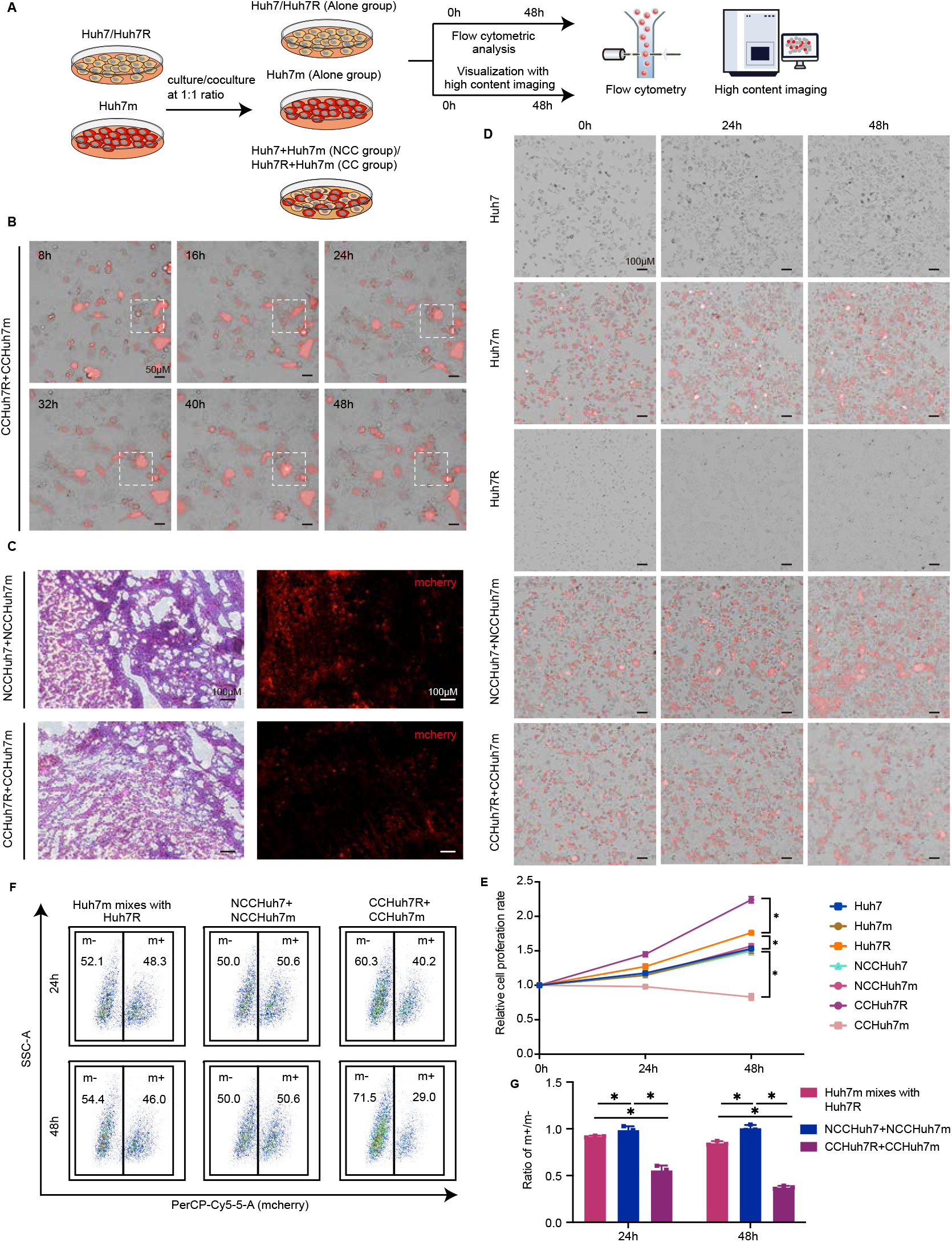
Cell competition drives the dominant growth of lenvatinib-resistant cells in HCC. (A) Schematic workflow of our experimental strategy. Huh7 or Huh7R was cocultured with Huh7m at 1:1 ratio or cultured alone respectively. Cells were cultured for 48h after cell attachment and then performed by high content imaging analysis and flow cytometry analysis. (B) High content imaging of cells growth over a time course of coculture after cells attachment in CC group (CCHuh7R+CCHuh7m) using 20X objective lens (Details see Video 5). (C) H&E stain and subsequent fluorescence imaging of cells in NCC group- and CC group-mice. (D) High content imaging of cells growth at 0h, 24h, 48h after cell attachment in Alone group (Huh7, Huh7m, Huh7R), NCC group (NCCHuh7+NCCHuh7m) and CC group using 10X objective lens (Details see Video 6-10). (E) Quantitative statistics of cell proliferation rate in (D). (F) Flow cytometry analysis of cell proportion in NCC group, CC group and single Huh7m mixed with single Huh7R at 24h and 48h after cell attachment. mcherry+ (m+): Huh7m, NCCHuh7m, CCHuh7m; mcherry- (m-): Huh7R, NCCHuh7, CCHuh7R. (G) Quantitative statistics of ratio (m+/m-) in (F). Three independent experiments were conducted, and the values are represented by means ± SD using two-way ANOVA with Turkey’s multiple comparisons test. *p < 0.05, ns = non-statistically significant. See also Figure S1.

Together, these results suggest that there exists cell competition between lenvatinib-resistant HCC cells and sensitive HCC cells under coculture condition and importantly the cell competition behavior further endows HCC lenvatinib-resistant cell with a growth advantage at the price of lenvatinib-sensitive cell.

### Lenvatinib-resistant cells present transcriptionally upregulated glycolytic metabolism in cell competition condition

To uncover the biological mechanism behind the phenomenon of cell competition between lenvatinib-resistant HCC cells and sensitive HCC cells, cocultured cells including CCHuh7R\CCHuh7m and NCCHuh7\NCCHuh7m were separated by flow cytometry cell sorting and then used for RNA sequencing (RNA-seq) and bioinformatics analysis; meanwhile alone cultured Huh7, Huh7m and Huh7R were subjected to the same processing (**Fig 2A**). According to GSEA analysis based on the results of differential expression analysis (**Fig 2B-E**), oxidative phosphorylation was significantly down-regulated in CCHuh7R compared to CCHuh7m cells (**Fig 2F**). Of note, similar results of mitochondria mass or oxidative phosphorylation related activities (mitochondrial respiratory chain complex assembly, mitochondrial gene expression, mitochondrial transmembrane transport, mitochondrial translation, mitochondrial RNA metabolic process, oxidative phosphorylation) was also found in CCHuh7R vs NCCHuh7/CCHuh7m and Huh7R vs Huh7 (**Fig EV2A-F**). Combining with the tendency of growth rate as described previously (**Fig 1D and E; Video6-10**) and the demonstration that decreased oxidative phosphorylation activity is accompanied by an increase in glycolytic flux and the switch of glycolysis (de la Cova *et al*, 2014) which is a hallmark of the increased proliferation of cancer cell (Hanahan & Weinberg, 2011), we infer that there exists a significantly differential energy metabolism activity or metabolic switch contributing to cell competition behavior. Consistent with our hypothesis, further GSEA analysis manifested that glycolysis pathway was significantly enriched and up-regulated in CCHuh7R compared to CCHuh7m cells, NCCHuh7 or Huh7R; meanwhile, no significant metabolism-related pathways were found in NCCHuh7 vs NCCHuh7m/Huh7m/Huh7 (**Fig 2G-I**). Moreover, considerable evidence implicates the occurrence of energy metabolism changes in cell competition: Myc upregulation (Winner) exhibits the increased glucose uptake and glycolysis activity in Drosophila imaginal plate (de la Cova *et al*., 2014); metabolic processes are regulatable and changed by transcriptional regulators related to cell competition (Lawlor *et al*, 2020). The above results and the previous studies support that energy metabolism reprogramming may be involved in the cell competition between lenvatinib-resistant cells and lenvatinib-sensitive cells in HCC.

**Figure 2.**
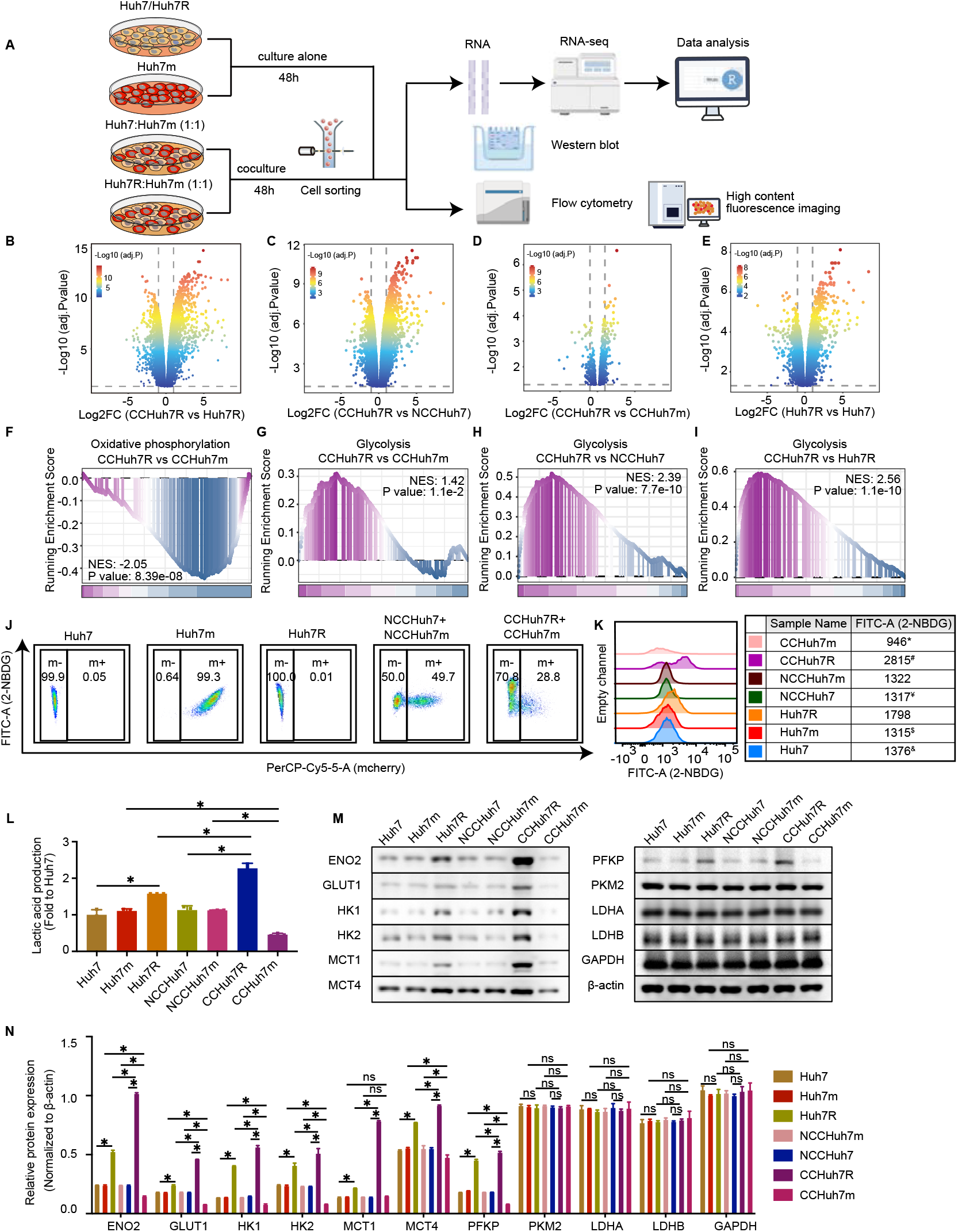
Lenvatinib-resistant cells display upregulated glycolytic metabolism in cell competition at transcriptomic level. (A) Schematic workflow of our experimental strategy by Figdraw. Huh7, Huh7m and Huh7R cultured alone; Huh7 or Huh7R cocultured with Huh7m at 1:1 ratio respectively. Cells were cultured for 48h after cell attachment and then coculture groups were separated using flow cytometry sorting for RNA-seq, Western blot and other purposes. (B) Volcano plot showing differentially expressed genes in CCHuh7R vs CCHuh7R group. (C) As in (B) but in CCHuh7R vs NCCHuh7 group. (D) As in (B) but in CCHuh7R vs CCHuh7m group. (E) As in (B) but in Huh7R vs Huh7 group. (F) GSEA analysis of oxidative phosphorylation enrichment in CCHuh7R vs CCHuh7m group. (G) GSEA analysis of glycolysis enrichment in CCHuh7R vs CCHuh7m group. (H) As in (G) but in CCHuh7R vs NCCHuh7 group. (I) As in (G) but in CCHuh7R vs Huh7R group. (J). Flow cytometry analysis of 2-NBDG-FITC-A and cell proportion at 48h after cell attachment in Alone group, NCC group and CC group. (K) Mountain map of 2-NBDG-FITC-A in (J). * : CCHuh7m vs NCCHuh7m (p < 0.05); $ : CCHuh7m vs Huh7m (p < 0.05); ¥ : CCHuh7R vs NCCHuh7 (p < 0.05); # : CCHuh7R vs Huh7R (p < 0.05); & : Huh7R vs Huh7 (p < 0.05). (L) Cellular lactic acid production levels at 48h after cell attachment in Alone group, NCC group and CC group by flow cytometry cell sorting. (M) Protein levels of ENO2, GLUT1, HK1, HK2, MCT1, MCT4, PFKP, PKM2, LDHA, LDHB and GAPDH at 48h after cell attachment in Alone group, NCC group and CC group detected by western blot. (N) Quantitative statistics of protein levels in (M). Three independent experiments were conducted, and the values are represented by means ± SD using two-way ANOVA with Turkey’s multiple comparisons test *p < 0.05, ns = non-statistically significant. See also Figure S2.

To further confirm the findings from RNA-seq analysis, cellular functional experiments on energy metabolism were performed in this study. As a marker for glycolysis, the indigestible glucose analogue 2-NBDG (Fantin *et al*, 2006; Intlekofer & Finley, 2019) was used to detect glucose uptake in indicated groups. Notably, in the cell competition environment, CCHuh7R with a significantly major population dominance displayed higher 2-NBDG level than Huh7R/NCCHuh7, observed in flow cytometry analysis; in contrast, the significant decreased 2-NBDG level was observed in CCHuh7m vs Huh7m/NCCHuh7m (**Fig 2J and K**). Additionally, to validate the integrity of glycolysis process, the end-product lactic acid was used to determine glycolytic activity. Consistent with the 2-NBDG results, the level of cellular lactic acid was sigficantly higher in CCHuh7R than in Huh7R/NCCHuh7 but relatively lower in CCHuh7m than in Huh7m/NCCHuh7m (**Fig 2L**). Considering substrates and glycolysis metabolic activity was boosted, partial glycolysis-related key metabolic proteins including ENO2, GLUT1, HK1, HK2, MCT1, MCT4, PFKP were detected and found to be highly expressed in CCHuh7R compared to Huh7R/NCCHuh7 cells; spontaneously, as the loser in competitive environment, CCHuh7m displayed a lower expression of glycolysis-related metabolic proteins compared to Huh7m/NCCHuh7m cells (**Fig 2M and N**). These results confirmed the hints from our bioinformatic analysis. Collectively, it indicates that enhanced glycolysis of lenvatinib-resistant cells may contribute to its outcompeting against sensitive cells in HCC.

It is well-known that energy metabolism forms of mitochondrial oxidative phosphorylation and glycolysis are interconnected process (Fantin *et al*., 2006), and tumor cells obtain adequate nutrients for proliferation and better survival in the stressful microenvironment via shifting energy production from mitochondrial oxidative phosphorylation to glycolysis (Intlekofer & Finley, 2019; Schworer *et al*, 2019). Thus, to unreval the effects of cellular energy metabolism on cell competition between lenvatinib-resistant HCC cells and sensitive HCC cells, we first examined the mitochondrial membrane potential by using Rhodamine 123, a positively charged green fluorescent dye that aggregated in active mitochondria and determined by a membrane potential gradient through their inner membranes. The result showed that the mitochondrial membrane potential was significantly weakened in Huh7R compared to Huh7 cells, suggesting the reduced level of oxidative phosphorylation, and CCHuh7R exhibited significantly lower mitochondrial membrane potential than Huh7R/NCCHuh7; moreover, the mitochondrial membrane potential was also remarkably decreased in CCHuh7m compared to those in Huh7m/NCCHuh7m cells, potentially underlying the incompetent status of CCHuh7m as a loser, though its oxidative phosphorylation levels was higher than CCHuh7R (**Fig EV2G-H**). Furthermore, Mito-Tracker Green, a mitochondrial morphology indicator, was used and it was found that energy metabolism shift significantly observed in the cell competition model and decreased mitochondrial mass of CCHuh7R compared with NCCHu7/Huh7R (**Fig EV2I-K**). Of note, the mitochondria of CCHuh7R displayed morphologically grainy and was exacerbated with a time dependent manner(**Fig EV2L)**, which was constent with the mitochondria morphology of mitophagy process (Dorn & Kitsis, 2015). Further, lower expression levels of IDH3A, COX7A, SDHA, UQCRC2, OGDH and CS proteins, which are rate-limiting enzymes in oxidative phosphorylation, were displayed in CCHuh7R compared to Huh7R/NCCHuh7 cells; while CCHuh7m exhibited significantly higher expression levels of those proteins than Huh7m/ NCCHuh7m (**Fig EV2M and N**).

Altogether, all these findings suggested that energy metabolism reprogramming may serve as a trigger of cell competition, in which mitochondrial oxidative phosphorylation is attenuated while glycolysis is enhanced in resistant cells, thereby driving lenvatinib resistance in HCC.

### Increased glycolytic flux supports the winner status of lenvatinib-resistant cells

Claire et al. reported that glucose availability in the microenvironment may also affect winner cells’ glycolysis activity though cell competition considerably improves winner cells’ glucose absorption which is modulated by transcription factors such as MYC and P53 (de la Cova *et al*., 2014). Although the experiments above suggested that contribution of energy metabolism reprogramming to cell competition between CCHuh7R and CCHuh7m, the impact of energy metabolism on cell competition behavior or which energy metabolism functioning as a vital role is unclear. Thus, we first investigated the effect of low glucose medium, which can inhibit glycolysis activity in HCC cells (Wu *et al*, 2021; Zappasodi *et al*, 2021) and afterwards observed functional indicators mentioned previously. Compared with the untreated group, low glucose-treated CCHuh7R reasonably appeared decreased 2-NBDG uptake but showed no difference in mitochondrial membrane potential, as observed by flow cytometry (**Fig 3A, B, D and E**). Notably, unaltered major population dominance of CCHuh7R (m+/m- < 1) compared with the untreated CCHuh7R (m+/m- < 1) indicated that cell competition phenomenon was weakened but not abolished (m+/m-: Low glucose group > Control group; p < 0.05) in low glucose condition (**Fig 3C**). On the other hand, to assess whether oxidative phosphorylation has similar effects on cell competition and the fitness of HCC lenvatinib-resistant cells, we used an oxidative phosphorylation agonist reagent Calcitriol (1,25-Dihydroxyvitamin D3) (Ferreira *et al*, 2015; Zhang *et al*, 2021) and found that 2-NBDG uptake in CCHuh7R with Calcitriol treatment was unaffected compared with untreated CCHuh7R (**Fig 3A and B**). Interestingly, the treatment of Calcitriol not only significantly increased mitochondrial membrane potential of CCHuh7R (**Fig 3D and E**) but also aggravated its major population status when compared with the untreated one (m+/m- ratio; Calcitriol group<Control group: p<0.05), indicating that cell competition behaviour was further enhanced (**Fig 3F**). These results preliminarily suggest that both the energy restriction such as low glucose treatment and the regulation of mitochondria oxidative phosphorylation metabolic levels may have a considerable but opposite impact on cell competition.

**Figure 3.**
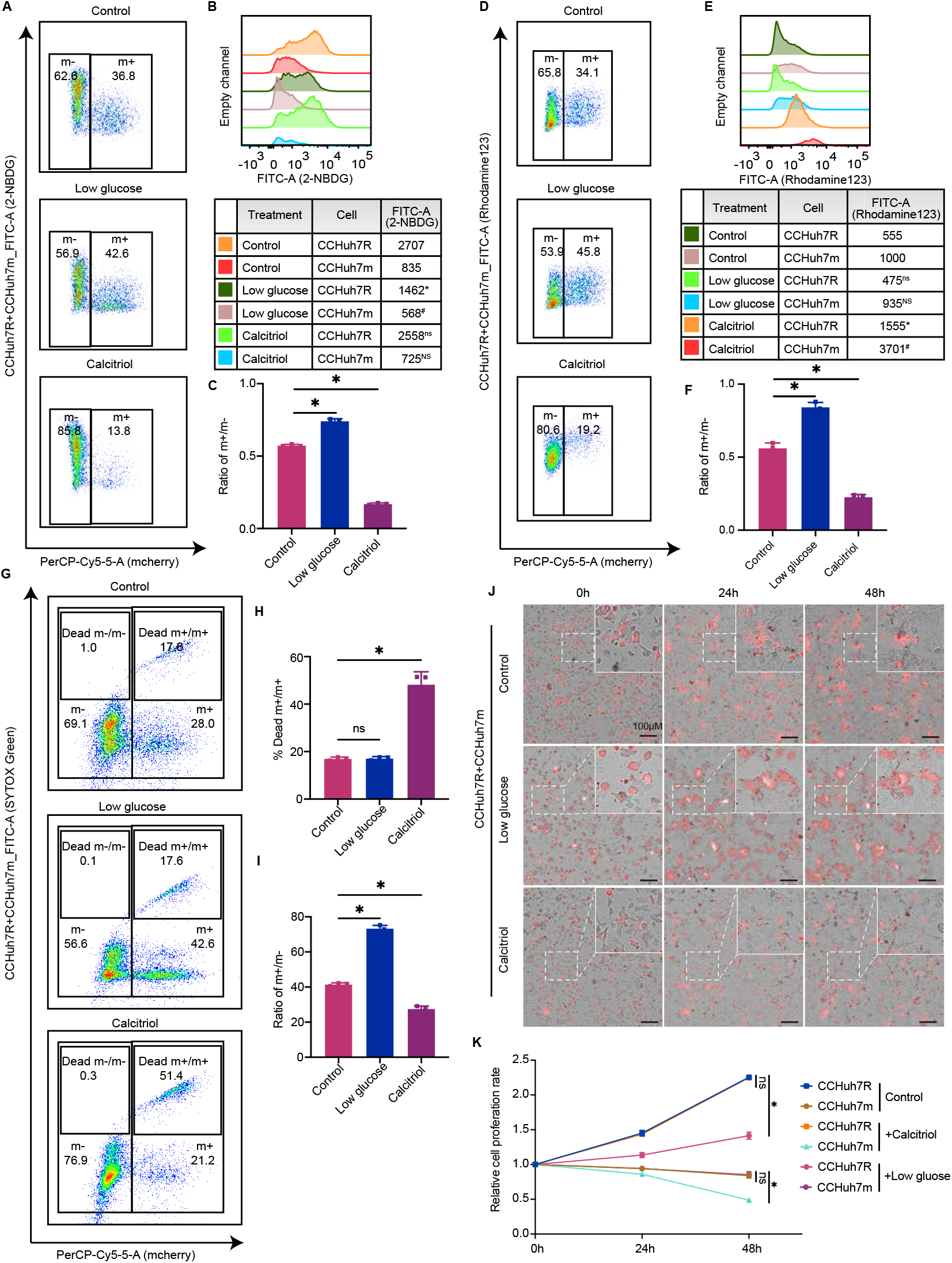
Enhanced glycolysis supports the winner status of lenvatinib-resistant cells. (A) Flow cytometry analysis of 2-NBDG-FITC-A and cell proportion at 48h after cell attachment in CC group with low glucose or Calcitriol treatment. (B) Quantitative statistics of 2-NBDG-FITC-A intensity in (A). * : CCHuh7R (Low glucose) vs CCHuh7R (Control) (p < 0.05); # : CCHuh7m (Low glucose) vs CCHuh7m (Control) (p < 0.05); ns : CCHuh7R (Calcitriol) vs CCHuh7R (Control) (p > 0.05); NS : CCHuh7m (Calcitriol) vs CCHuh7m (Control) (p > 0.05). (C) Quantitative statistics of ratio of m+/m- in (A). (D) Flow cytometry analysis of mitochondrial membrane potential (Rhodamine123-FITC-A) and cell proportion at 48h after cell attachment in CC group with low glucose or Calcitriol treatment. (E) Quantitative statistics of 2-NBDG-FITC-A in (D). ns : CCHuh7R (Low glucose) vs CCHuh7R (Control) (p > 0.05); NS : CCHuh7m (Low glucose) vs CCHuh7m (Control) (p > 0.05); * : CCHuh7R (Calcitriol) vs CCHuh7R (Control) (p > 0.05); # : CCHuh7m (Calcitriol) vs CCHuh7m (Control) (p > 0.05). (F) Quantitative statistics of ratio of m+/m- in (D). (G) Flow cytometry analysis of cell death (SYTOX Green-FITC-A) and cell proportion at 48h after cell attachment in CC group with low glucose or Calcitriol treatment. (H) Quantitative statistics of Dead m+/m+ ratio in (G). (I) Quantitative statistics of m+/m+ ratio in (G). (J) High content imaging of cells growth at 0, 24h, 48h after cell attachment in CC group with low glucose or Calcitriol treatment using 10X objective lens. (K) Quantitative statistics of cell number in (J) (Details see Video 11-13). Three independent experiments were conducted, and the values are represented by means ± SD using two-way ANOVA with Turkey’s multiple comparisons test (B, E and K) or Ordinary one-way ANOVA with Sidak’s multiple comparisons test (C, F, H and I). *p < 0.05, ns = non-statistically significant.

To further identify that the detailed effects of energy metabolism reprogramming on cell competition in HCC, we performed a SYTOX Green assay to observe the relative survival state of resistant HCC cells to sensitive HCC cells and compared the death ratio of CCHuh7m/CCHuh7R with specific treatments in a cell competition scenario. Noticeably, low glucose treatment had no significant effect on the death ratios of CCHuh7m (Dead CCHuh7m vs CCHuh7m) and CCHuh7R (Dead CCHuh7R vs CCHuh7R), but the ratio of CCHuh7m vs CCHuh7R remarkably increased. However, upon Calcitriol treatment, more than 2-fold was observed in the death ratio of CCHuh7m, accompanied by unaltered CCHuh7R deaths and reduced m+/m- (**Fig 3G-I**). These results imply that the variation of cell competition between CCHuh7R and CCHuh7m most likely depends on their distinct metabolic features. Thus, we then examined and compared the growth rates of CCHuh7R and CCHuh7m under the above additions in a setting of cell competition. As expected, low glucose treatment mainly attenuated the growth of CCHuh7R but did not strongly suppress the growth of CCHuh7m. Nevertheless, Calcitriol could profoundly reduce CCHuh7m viability but had no notable effect on the growth of CCHuh7R (**Fig 3J and K; Video 11-13**). These results suggested, energy restriction contributed to the increase of the CCHuh7m/CCHuh7R ratio by inhibiting CCHuh7R growth, whereas the activated oxidative phosphorylation levels led to the diminishment of the CCHuh7m/CCHuh7R ratio by increasing CCHuh7m death.

Collectively, these in vitro studies above prove that enhanced glycolysis, but not oxidative phosphorylation, is the principal competitive strategy by maintaining the winner status of HCC lenvatinib-resistant cells.

### BNIP3 mediated mitophagy is required for enhanced glycolysis of lenvatinib-resistant cells (winner)

To further clarify the molecular mechanism by which enhanced glycolysis activity of CCHuh7R contributing to its cell competition advantage in our model, we analyzed the differentially expressed genes (DEGs) in CCHuh7R vs Huh7R group and CCHuh7R vs NCCHuh7 group. UpSet plot analysis identified twelve overlapping genes in Top 50 DEGs of each group which were ranked by FDR (**Fig 4A**), demonstrating that these genes’ expressions were remarkably changed in cell competition scenario. Interestingly, BNIP3, a marker of mitophagy occurrence and its participation in improving glycolysis activities including glucose uptake and lactic acid production (O’Sullivan *et al*, 2015; Springer *et al*, 2021; Thomas *et al*, 2011), exhibited significant up-regulation (**Fig 4A**). Considering the findings in attenuated mitochondrial oxidative phosphorylation in CCHuh7R validated by GSEA analysis and cellular functional experiments, we speculated that cell competition may heighten glycolytic flux of HCC lenvatinib-resistant cells through BNIP3- mediated mitophagy process for eliminating superfluous or aberrant mitochondria, which is the primary cause of the reduced mitochondrial oxidative phosphorylation and membrane potential. To validate this hypothesis, a series of bioinformatic analyses and relevant experiments were performed. As expected, CCHuh7R showed significant down-regulated mitochondrial mass/activities by GSVA analysis and differential expression analysis based on customized gene set of GO and, surprisingly, two mitophagy-related activities (positive regulation of mitophagy in response to mitochondrial depolarization, regulation of autophagy of mitochondrion in response to mitochondrial depolarization) were notably enriched in these activities (**Fig EV3A and B**). Furthermore, the identification of colocalization between Cy5.5-LC3B-positive autophagosome and TOMM20-tagged mitochondria detected by immunofluorescence showed that there was colocalization in CCHuh7R and Huh7R and it occurred significantly more in CCHuh7R than in Huh7R (**Fig 4B**); moreover, higher expression of mitophagy-related protein levels (LC3B, BNIP3) and lower expression of TOMM20 representing mitochondria mass were further observed in CCHuh7R (**Fig 4C and D**). The above results suggested that CCHuh7R indeed had more abnormal mitochondria and experienced more mitophagy than Huh7R, which was also observed in HCC sorafenib resistance (Chen *et al*, 2019; Wu *et al*, 2020). Subsequently, we specifically focused on the mitophagy pathway, and found that BNIP3 with the minimum FDR displayed remarkable expression difference in two comparative groups as showed by heatmap (**Fig EV3C and D**). These data again substantiated our hypothesis. Although we have identified the occurrence of mitochondrial aberration and mitophagy of HCC lenvatinib-resistant cell in cell competition scenario, the correlation between BNIP3-mediated mitophagy and glycolysis was still not completely clear. Therefore, clinical HCC specimens in TCGA and ICGC database (TCGA-LIHC chort and ICGC-LIHC chort) were analysed and categorized into BNIP3 high- and low-group according to the median value of BNIP3 expression. GSEA analyses from both cohorts showed that glycolysis pathway was remarkably enriched and up-regulated in high expression of BNIP3 group and had a positive correlation with BNIP3 expression based on ssGSEA analysis (**Fig 4E and F, Fig EV3E and F**).

**Figure 4.**
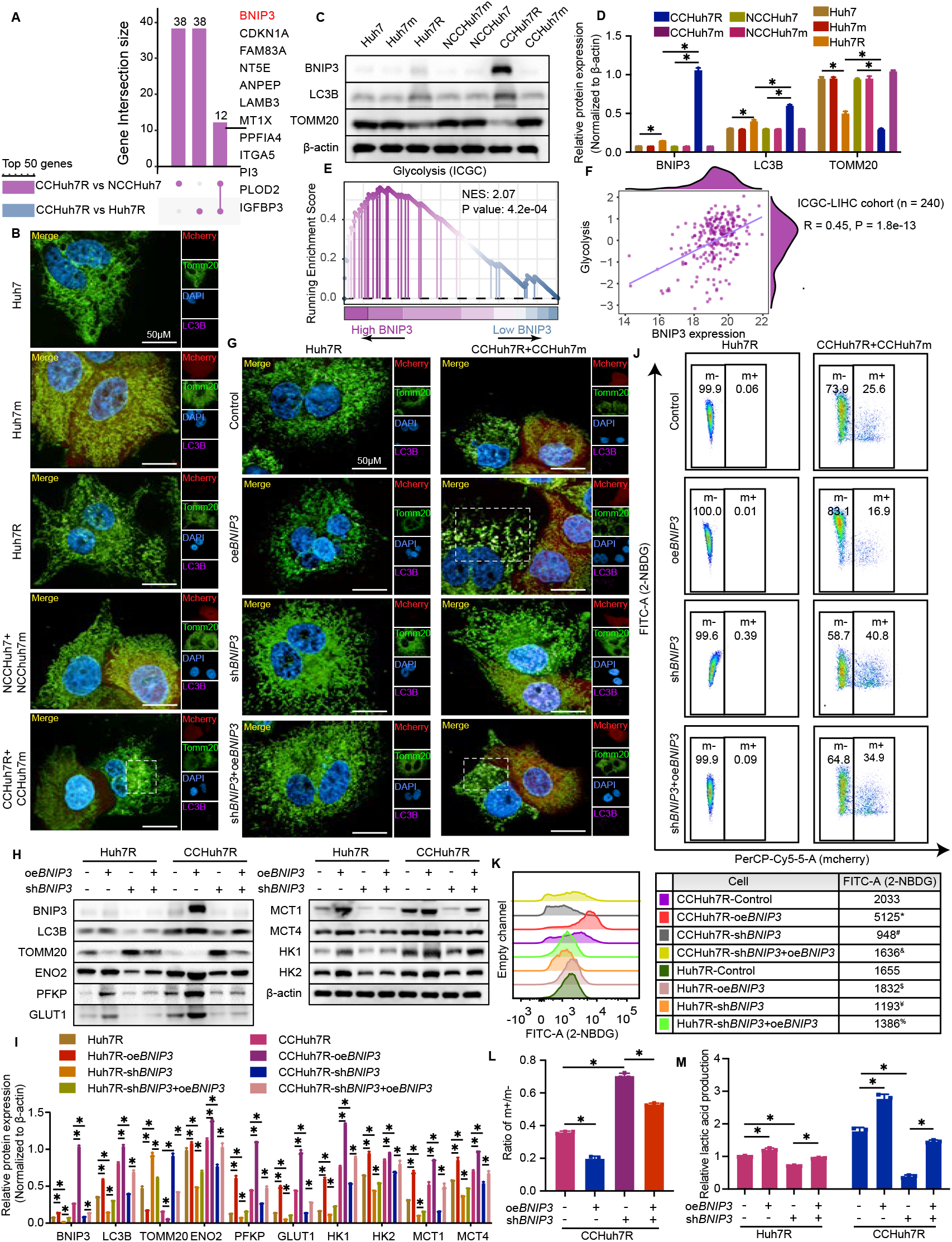
Enhanced glycolytic flux of lenvatinib-resistant cells (winner) requires BNIP3 mediated mitophagy. (A) UpSet plot showing Top 50 DEGs’ overlapping genes in CCHuh7R vs Huh7R group and CCHuh7R vs NCCHuh7 group. (B) High content immunofluorescence imaging of colocalization of autophagosomes (Cy5.5-LC3: purple) and mitochondria (TOMM20: green) at 48h after cell attachment in Alone group, NCC group and CC group using 63X water immersion objective lens. (C) Protein levels of LC3B, TOMM20 and BNIP3 at 48h after cell attachment in Alone group, NCC group and CC group detected by western blot. (D) Quantitative statistics of protein levels in (C). (E) GSEA analysis of glycolysis enrichment in BNIP3 high-expression vs BNIP3 low-expression group of ICGC-LIHC cohort. (F) Correlation analysis of BNIP3 expression and glycolysis pathway in ICGC-LIHC cohort. (G) High content immunofluorescence imaging of colocalization of autophagosomes (Cy5.5-LC3: purple) and mitochondria (TOMM20: green) at 48h after cell attachment in Huh7R (Alone group) /CCHuh7R (CC group) transfected with shRNA against *BNIP3* (sh*BNIP3*), *BNIP3* overexpression plasmid (oe*BNIP3*) or oe*BNIP3*+sh*BNIP3* using 63X water immersion objective lens. (H) Protein levels of LC3B, TOMM20, BNIP3, ENO2, GLUT1, HK1, HK2, MCT1, MCT4 and PFKP at 48h after cell attachment in Huh7R (Alone group)/CCHuh7R (CC group) transfected with sh*BNIP3*, oe*BNIP3* or sh*BNIP3*+oe*BNIP3* detected by western blot. (I) Quantitative statistics of protein levels in (H). (J) Flow cytometry analysis of 2-NBDG-FITC-A and cell proportion at 48h after cell attachment in Huh7R (Alone group) /CCHuh7R (CC group) transfected with sh*BNIP3*, oe*BNIP3* or sh*BNIP3*+oe*BNIP3*. (K) Mountain map of 2-NBDG-FITC-A in (J). * : CCHuh7R-oe*BNIP3* vs CCHuh7R-Control (p < 0.05); # : CCHuh7R-sh*BNIP3* vs CCHuh7R-Control (p < 0.05); & : CCHuh7R-sh*BNIP3+*oe*BNIP3* vs CCHuh7R-sh*BNIP3* (p < 0.05); $ : Huh7R-oe*BNIP3* vs Huh7R-Control (p < 0.05); ¥ : Huh7R-sh*BNIP3* vs Huh7R-Control (p < 0.05); % : Huh7R-sh*BNIP3+*oe*BNIP3* vs Huh7R-sh*BNIP3* (p < 0.05). (L) Quantitative statistics of ratio m+/m- in (J). (M) Cellular lactic acid production levels at 48h after cell attachment in Huh7R (Alone group) /CCHuh7R (CC group) transfected with sh*BNIP3*, oe*BNIP3* or sh*BNIP3*+oe*BNIP3* by flow cytometry cell sorting. Three independent experiments were conducted, and the values are represented by means ± SD using two-way ANOVA with multiple comparisons test (D, I and M) or Ordinary one-way ANOVA with Sidak’s multiple comparisons test (L). *p < 0.05, ns = non-statistically significant.

To further prove the finding above from bioinformatic analyses, we constructed genetically modified Huh7R overexpressing *BNIP3* (oe*BNIP3*) or lacking *BNIP3* (sh*BNIP3*) with or without the coculture of Huh7m (Huh7R-sh*BNIP3*, Huh7R-oe*BNIP3*, CCHuh7R-sh*BNIP3*, CCHuh7R-oe*BNIP3*). Considering the rigor and logicality of validation, metabolic indicators of modified Huh7R/CCHuh7R were also investigated under another experimental manipulation where the expression of BNIP3 protein was induced in a mosaic form by transient expression (Huh7R-sh*BNIP3*+oe*BNIP3*, CCHuh7R-sh*BNIP3*+oe*BNIP3*). We found that the knockdown of *BNIP3* significantly inhibited the colocalization of Cy5.5-LC3B and TOMM20 in CCHuh7R cells(**Fig 4G**). Furthermore, CCHuh7R-sh*BNIP3* relative to Huh7R-sh*BNIP3* expressed higher TOMM20, which could be rescued by the transient transfection of *BNIP3*. Conversely, *BNIP3* overexpression achieved opposite effects (**Fig 4H and I**). These results indicated the importance of BNIP3 which induces the occurrence of mitophagy for eliminating abnormal mitochondria in CCHuh7R, although there existed a moderate change in Huh7R in line with what has been reported previously (Subramanian *et al*, 2021). Noteworthily and importantly, lacking *BNIP3* decreased the expression levels of glycolysis-related proteins (ENO2, GLUT1, HK1, HK2, MCT1, MCT4, PFKP) and 2-NBDG and lactic acid productionin in Huh7R and CCHuh7R (**Fig 4H, I, J, K and M**). However, this manipulation displayed positive but limited alteration of the competitive advantage of CCHuh7R (**Fig 4J and L**). Combining with the contrary results of CCHuh7R-oe*BNIP3*, these results provide evidence that BNIP3 could significantly modulate glycolysis of CCHuh7R but not change the decisive trend of cell competition between lenvatinib-resistant HCC cells and sensitive HCC cells.

In summary, we conclude that lenvatinib-resistant HCC cells hold the increased glycolytic flux through BNIP3-mediated mitophagy, thereby maintaining its winner status.

### BNIP3-regulated ENO2 orchestrates the effect of mitophagy on glycolysis of lenvatinib-resistant cells (winner)

Next, we sought to uncover the mechanism how mitophagy regulates glycolysis or whether there exists major glycolysis target molecular regulated by mitophagy. In the previous western blot results, we noticed that ENO2, a key enzyme of glycolysis process, showed a distinct variation in malignant phenotype-related glycolysis metabolism of CCHuh7R (**Fig 2M and N**, **Fig 4H and I**), verified by other cancers (Kim *et al*, 2019; Sun *et al*, 2021; Zheng *et al*, 2020). Intriguingly, specific glycolysis pathway analysis also showed that ENO2 with the minimum FDR displayed the biggest expression difference in CCHuh7R vs Huh7R group and CCHuh7R vs NCCHuh7 group depicted by heatmap and UpSet plot (**Fig 5A-C**). The above results potentially implied that ENO2 may be a critical effector in competitive cascade reaction of CCHuh7R. Therefore, we genetically test whether ENO2 exerts specific effect on glycolysis activity and competitive behaviour of HCC Lenvatinib-resistant cell. Strikingly, we found that *ENO2* deficiency (sh*ENO2*) specifically contributed to dramatic decrease in 2-NBDG (**Fig 5D and E**) and lactic acid production of CCHuh7R (CCHuh7R-sh*ENO2*) in cell competition scenario, which could be abrogated by the transient supplement of ENO2 (CCHuh7R-sh*ENO2*+oe*ENO2*); corresponding changes could be observed in the administration of ENO2 overexpression (CCHuh7R-oe*ENO2*) (**Fig 5G**). Nevertheless, both Huh7R and CCHuh7R exhibited no remarkable change in glycolysis and mitophagy-related proteins expression (**Fig 5H and I**) and the colocalization between Cy5.5-LC3B and TOMM20 (**Fig 5J**), no matter which genetic modification was performed (sh*ENO2*, oe*ENO2*). These results represented ENO2 functions as the unique role in enhanced glycolysis activity of CCHuh7R, likely as a downstream of BNIP3. Of note, the ratio of m+/m- obtained a marked but limited increase even in the case of the reduction of ENO2-represented glycolysis levels of CCHuh7R (**Fig 5F**), potentially manifesting that the variation of ENO2 expression is still not efficient enough to eliminate cell competition phenomenon. To elucidate the detailed relationship between mitophagy, glycolysis and ENO2, we overexpressed BNIP3 in *ENO2*-deficient Huh7R with or without coculture with Huh7m (Huh7R-sh*ENO2*+oe*BNIP3*, CCHuh7R-sh*ENO2*+oe*BNIP3*). As expected, *BNIP3*-temporary overexpression rescued the detected effects of CCHuh7R- sh*ENO2*, which was reasonably inferior to CCHuh7R-oe*BNIP3* (**Fig EV4A-F**). These results suggested that there existed BNIP3-ENO2 regulatory axis which featured in enhanced glycolysis of CCHuh7R in cell competition scenario.

**Figure 5.**
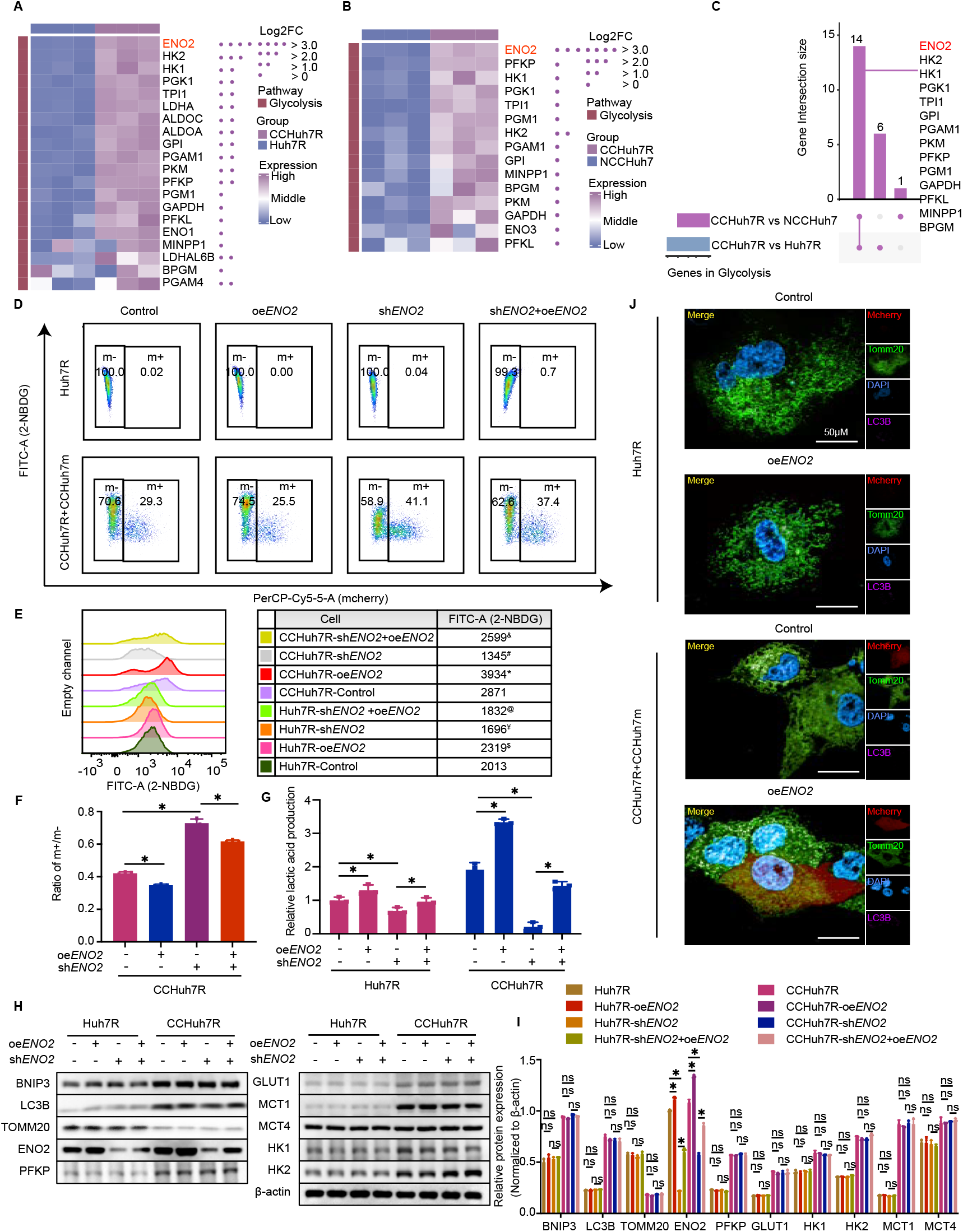
The effect of glycolysis of winner is ENO2 dependent. (A) Heatmap exhibiting expression levels of glycolysis-related genes between CCHuh7R and Huh7R. (B) As in (A) but between CCHuh7R and NCCHuh7. (C) UpSet plot displaying overlapping glycolysis-related genes based on (A) and (B). (D) Flow cytometry analysis of 2-NBDG-FITC-A and cell proportion at 48h after cell attachment in Huh7R (Alone group) /CCHuh7R (CC group) transfected with shRNA against *ENO2* (sh*ENO2*), *ENO2* overexpression plasmid (oe*ENO2*) or oe*ENO2*+sh*ENO2*. (E) Mountain map of 2-NBDG-FITC-A in (D). * : CCHuh7R-oe*ENO2* vs CCHuh7R-Control (p < 0.05); # : CCHuh7R-sh*ENO2* vs CCHuh7R-Control (p < 0.05); & : CCHuh7R-sh*ENO2+*oe*ENO2* vs CCHuh7R-sh*ENO2* (p < 0.05); $ : Huh7R-oe*ENO2* vs Huh7R-Control (p < 0.05); ¥ : Huh7R-sh*ENO2* vs Huh7R-Control (p < 0.05); % : Huh7R-sh*ENO2+*oe*ENO2* vs Huh7R-sh*ENO2* (p < 0.05). (F) Quantitative statistics of ratio m+/m- in (D). (G) Cellular lactic acid production levels at 48h after cell attachment in Huh7R (Alone group) /CCHuh7R (CC group) transfected with sh*ENO2*, oe*ENO2* or sh*ENO2*+oe*ENO2* by flow cytometry cell sorting. (H) Protein levels of LC3B, TOMM20, BNIP3, ENO2, GLUT1, HK1, HK2, MCT1, MCT4 and PFKP at 48h after cell attachment in Huh7R (Alone group) /CCHuh7R (CC group) transfected with sh*ENO2*, oe*ENO2* or sh*ENO2*+oe*ENO2* detected by western blot. (I) Quantitative statistics of protein levels in (H). (J) High content immunofluorescence imaging of colocalization of autophagosomes (Cy5.5-LC3: purple) and mitochondria (TOMM20: green) at 48h after cell attachment in Huh7R (Alone group) /CCHuh7R (CC group) transfected with sh*ENO2*, oe*ENO2* or sh*ENO2*+oe*ENO2* using 63X water immersion objective lens. Three independent experiments were conducted, and the values are represented by means ± SD using a two-way ANOVA with Turkey’s multiple comparisons test (G and I) or Ordinary one-way ANOVA with Sidak’s multiple comparisons test (F). *p < 0.05, ns = non-statistically significant. See also Figure S4.

Altogether, it implys that mitophagy strengthens glycolysis of HCC lenvatinib-resistant cells through BNIP3-ENO2 regulatory axis for constantly supporiting its competitive dominance.

### AMPK functions as a signaling bridge in BNIP3-ENO2 crosstalk in lenvatinib-resistant cells

In the above studies, we had confirmed that ENO2 is a specific target in multiple potential proteins of glycolysis which was heighten by BNIP3-mediated mitophagy in our cell competition model. However, in the stressful and nutritionally competitive environment, the logically biological relevance between resistant and sensitive HCC cells or whether there exists an inductor to convey the information caused by energy metabolism changes is unclear. Of note, besides glycolysis pathway, previous validation of GSEA analyses for confirming the relationship between BNIP3 and glycolysis pathway in two clinical cohorts also enriched up-regulated AMPK signaling pathway (**Fig 6A and B**), which participates in cellular energy sensing and provokes glucose uptake through AMPK heterotrimeric kinase complex (Sharma *et al*, 2008), as evident by western blot ananlysis (**Fig 6C and D**). In addition, ssGSEA analysis also showed that there existed a remarkably positive correlation between BNIP3 expression and AMPK signaling pathway or AMPK expression (catalytic subunit: PRKAA1, PRKAA2; non-catalytic subunit: PRKAB1) (**Fig 6E and F, Fig EV5A and B**). Hence, we speculated the BNIP3-AMPK-ENO2 crosstalk exists and functions as the regulator of glycolytic metabolism and promoter of competitive advantage phenotypes of lenvatinib-resistant HCC cells in cell competition scenario. Thus, we firstly set out to determine the effect of AMPK. Consistent with previous BNIP3 analyses, AMPK inhibition (AMPK-IN-3) contributed to the reduction of glycolysis levels and competitive dominance of CCHuh7R compared with control, which was rescued by the treatment of AMPK activator (AMPK-activator-2) (**Fig 6G-J**), providing the evidence that AMPK may serve as a inductor for modulating glycolysis and competitive dominance of CCHuh7R in cell competition scenario. Furthermore, we sought to systematically investigate the existence of the BNIP3-AMPK-ENO2 axis and its regulatory relationship in our competitive models. As expected, compared with corresponding control groups, activated AMPK not only increased ENO2 expression levels of CCHuh7R, but also attenuated the inhibitory effects of CCHuh7R-sh*ENO2* on the above glycolysis indicators (excepting unchanged AMPK protein expression level) and cell competition phenomenon (**Fig 6L and M, Fig EV5C, D, E and I**). Therefore, we summarized that AMPK-regulated glycolysis and competitive dominance of CCHuh7R requires the existance of ENO2. On the other hand, overexpressed BNIP3 up-regulated level of phosphor-AMPK protein while no significant alteration was found in the expression levels of BNIP3 and the colocalization of mitophagy-related proteins of CCHuh7R under AMPK inhibition or activation treatment (**Fig 6K, N and O**). Nevertheless, overexpressed BNIP3 up-regulated the level of phosphor-AMPK and elevated AMPK-IN-3-inhibited metabolic levels and enhanced the outcompeting of CCHuh7R in cell competition scenario (**Fig 6N and O, Fig EV5F, G, H and J**). Summarizing our current results and previous validation of BNIP3-ENO2 regulatory axis in CCHuh7R cells in cell competition scenario, we concluded that mitophagy-represented BNIP3 could regulate AMPK and ENO2 in CCHuh7R; more importantly, the effect of mitophagy-represented BNIP3 on glycolysis and competitive dominance phenotypes of CCHuh7R is AMPK signaling-dependent.

**Figure 6.**
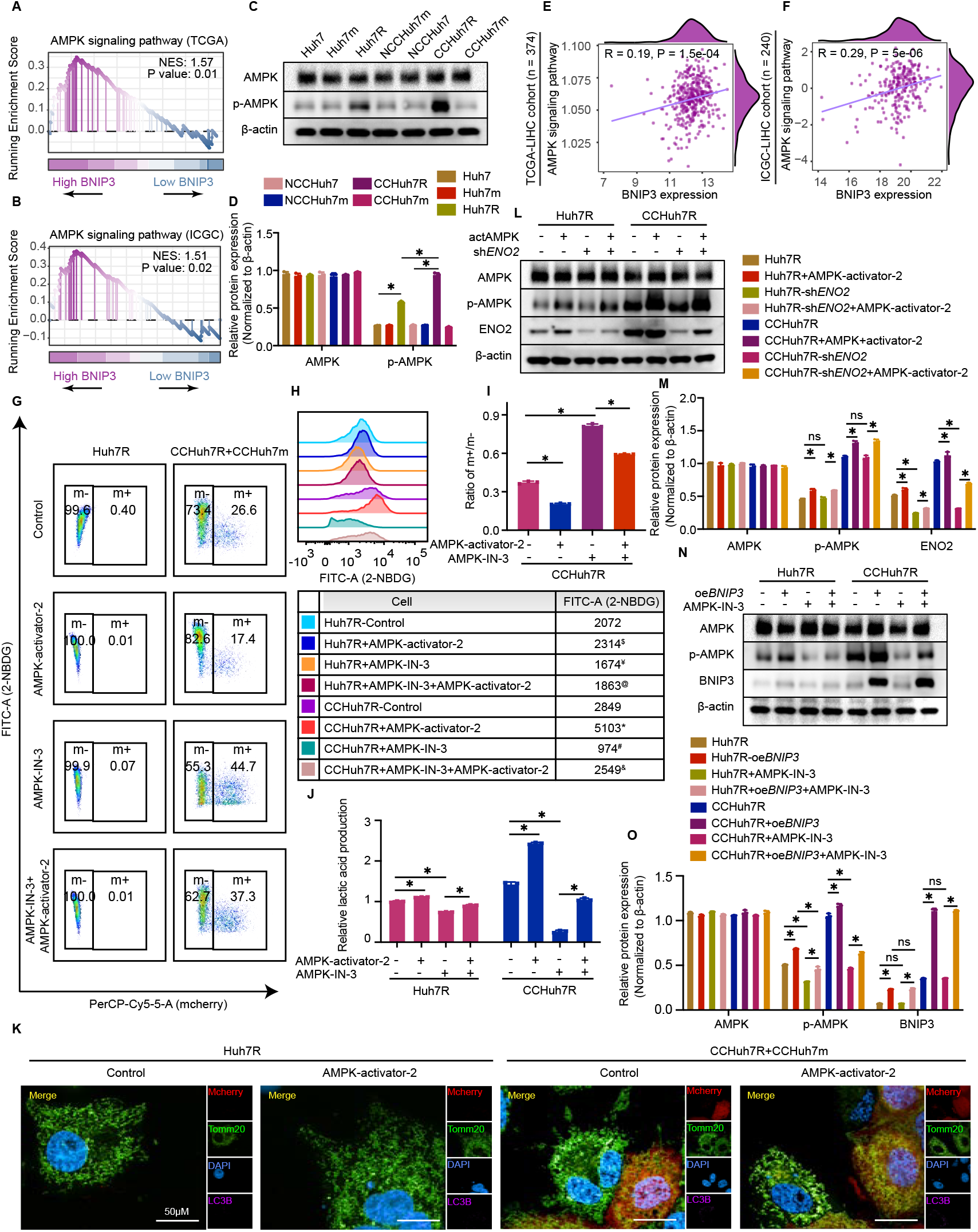
AMPK functions as a signaling bridge in BNIP3-ENO2 crosstalk in lenvatinib-resistant cells. (A) GSEA analysis of AMPK signaling pathway enrichment in BNIP3 high expression vs BNIP3 low expression group of TCGA-LIHC cohort. (B) As in (A) but in ICHC-LIHC corhort. (C) Protein levels of AMPK and p-AMPK at 48h after cell attachment in Alone group, NCC group and CC group detected by western blot. (D) Quantitative statistics of protein levels in (C). (E) Correlation analysis of BNIP3 expression and AMPK signaling pathway in TCGA-LIHC cohort. (F) As in (E) but in ICHC-LIHC corhort. (G) Flow cytometry analysis of 2-NBDG-FITC-A and cell proportion at 48h after cell attachment in Huh7R (Alone group) /CCHuh7R (CC group) treated with AMPK-activator-2, AMPK-IN-3 or AMPK-IN-3+AMPK-activator-2. (H) Mountain map of 2-NBDG-FITC-A in (G). * : CCHuh7R+AMPK-activator-2 vs CCHuh7R-Control (p < 0.05); # : CCHuh7R-AMPK-IN-3 vs CCHuh7R-Control (p < 0.05); & : CCHuh7R+AMPK-IN-3*+*AMPK-activator-2 vs CCHuh7R+AMPK-IN-3 (p < 0.05); $ : Huh7R-AMPK-activator-2 vs Huh7R-Control (p < 0.05); ¥ : Huh7R+AMPK-IN-3 vs Huh7R-Control (p < 0.05); % : Huh7R+AMPK-IN-3*+*AMPK-activator-2 vs Huh7R+AMPK-IN-3 (p < 0.05). (I) Quantitative statistics of ratio m+/m- in (G). (J) Cellular lactic acid production levels at 48h after cell attachment in Huh7R (Alone group) /CCHuh7R (CC group) treated with AMPK-activator-2, AMPK-IN-3 or AMPK-IN-3+AMPK-activator-2 by flow cytometry cell sorting. (K) High content immunofluorescence imaging of colocalization of autophagosomes (Cy5.5-LC3: purple) and mitochondria (TOMM20: green) at 48h after cell attachment in Huh7R (Alone group) /CCHuh7R (CC group) treated with AMPK-activator-2 using 63X water immersion objective lens. (L) Protein levels of AMPK, p-AMPK and ENO2 at 48h after cell attachment in Huh7R (Alone group) /CCHuh7R (CC group) treated with AMPK-activator-2, sh*ENO2* or sh*ENO2*+AMPK-activator-2 detected by western blot. (M) Quantitative statistics of protein levels in (L). (N) Protein levels of AMPK, p-AMPK and BNIP3 at 48h after cell attachment in Huh7R (Alone group) /CCHuh7R (CC group) treated with oe*BNIP3*, AMPK-IN-3 or oe*BNIP3*+AMPK-IN-3 detected by western blot. (O) Quantitative statistics of protein levels in (N). Three independent experiments were conducted, and the values are represented by means ± SD using a two-way ANOVA with Turkey’s multiple comparisons test (D, J, M and O) or Ordinary one-way ANOVA with Sidak’s multiple comparisons test (I). *p < 0.05, ns = non-statistically significant. See also Figure S5.

Totally, our accumulated summaries and evidence suggest that mitophagy facilitates glycolysis via BNIP3-AMPK-ENO2 crosstalk for persistently maintaining HCC lenvatinib-resistant cells’ competitive dominance.

## Discussion

Despite the advances in targeted drugs and treatment strategies for advanced HCC such as lenvatinib, cabozantinib, and multidrug combination strategy (Abou-Alfa *et al*, 2018; Finn *et al*, 2020; Kudo *et al*, 2018), clinical benefit remains dismal owing to the occurrence of acquired chemoresistance. At present, the molecular mechanism of chemoresistance in HCC has been explored and demonstrated thoroughly; the formulation of chemoresistance reversal therapies mainly relies on biological roots such as tumor burden and grow kinetics, tumor heterogeneity, tumor microenvironment and tumor immunity. Accordingly, treatment tactics including log kill method, synthetic lethal method and immunotherapies based on those biological foundations have been a research focus of drug-resistant therapy. Noticeably and unfortunately, the treatment methods based on tumor heterogeneity, a common phenomenon reported in tumors, are excluded from consideration. Tumor heterogeneity closely resembles a knotted Gordian Knot, where epigenetic changes and clonal evolution endow it with volatile phenotypes (Huang *et al*, 2020), demonstrating that strategies on account of tumor heterogeneity for drug resistance reversal are a challenge as well as a significantly promising approach. In recent years, increasing studies have reported that a heterogeneity-based biological phenotype, the supervision and exclusion of less suited “loser” cells by nearby “winner” cells, termed cell competition, is closely linked to tumorigenesis and development (Baker, 2020; Parker *et al*., 2021; Vishwakarma & Piddini, 2020). Importantly, a variety of cases and models have demonstrated that tumor cells with higher competence can constantly maintain and spread the most aggressive characteristics to outcompete relative less competent cells (Parker *et al*, 2020), which precisely conforms to the feature of tumor resistant cells (Gomes *et al*., 2022; Shen *et al*., 2013). Thus, these evidences suggest cell competition is expected to become a potential treatment basis.

In this study, through constructing mimetic lenvatinib-resistant cell and intercellular interaction system of advanced HCC, we first evidence that there exists cell competition between lenvatinib-resistant cells and lenvatinib-sensitive cells in HCC. During this process, lenvatinib-resistant cells obtain competitive dominance based on innate sub-foundation and further aggregate their proliferation characteristic at the price of lenvatinib-sensitive cells through energy metabolic shifting. It is reported that drug-resistant cells generally have superior capacities of proliferation, metastasis and metabolism (Gomes *et al*., 2022; Shen *et al*., 2013), which is in line with the verified characteristic of winner as well as the reason of its triumph (Di Gregorio *et al*, 2016; Eichenlaub *et al*, 2016). These results indicate the emergence of chemoresistance potentially is the result of survival of the fittest between resistant cells and primary cells during clinical treatments, and prove the study value of heterogeneity-mediated cell competition again. Nevertheless, the mechanism of cell competition between lenvatinib-resistant cells and lenvatinib-sensitive cells remain obscure.

Recent studies have manifested that activated glycolysis or energy metabolism differences are required to drive cell competition (Penzo-Mendez *et al*, 2015). Of note, transcriptome analyses show that under the cell competition environment, lenvatinib-resistant cells capture notably increased glycolysis activity but attenuated oxidative phosphorylation levels and decreased mitochondria mass; however, lenvatinib-sensitive cells obtain opposite metabolic features in cell competition. Importantly, results of in-vitro experiments including elevated glucose uptake and lactic acid secretion, high-expressed glycolysis key enzymes, low expression of rate-limiting enzymes in oxidative phosphorylation, reduced mitochondrial membrane potential and decreased mitochondrial mass have been observed and verified in lenvatinib-resistant cells; further, based on the illustration that glucose availability also has an effect upon winner’s glycolysis though cell competition considerably boosts winner’s glucose absorption, low glucose shows decreased glucose uptake and attenuated competitive dominance of lenvatinib-resistant cells through the influence on their growth; nevertheless, oxidative phosphorylation agonist Calcitriol has no effect on glucose uptake but notably exacerbates cell competition through promoting opponent’s death. These results imply that it is the metabolic differences between lenvatinib-resistant cells and lenvatinib-sensitive cells that lead to their distinct competition fitness. As we know, mitochondrial oxidative phosphorylation and glycolysis are crosslinked process (Fantin *et al*., 2006) and tumor cells grab enough nutrients for survival under various pressure through transforming into glycolysis following the Warburg effect (Intlekofer & Finley, 2019; Schworer *et al*., 2019). Accordingly, it is reasonable and easy to infer that lenvatinib-resistant cells competitively obtain sufficient glucose substrates for survival and continually maintain their winner status through intelligently switching energy metabolism from mitochondrial oxidative phosphorylation to principal glycolysis under the stressful cell competition environment. However, although decreased mitochondrial membrane potential caused by energy metabolism shifting is an appropriate way to maximize survival, the reason why that occurred and aggregated by Calcitriol in lenvatinib-sensitive cells, which is contrary to its fate remains unclear. It is possible that lenvatinib-sensitive cells confronting certain oxidative stress such as elevated ROS levels in this competitive environment (Kohashi *et al*, 2021; Mori *et al*, 2022). In addition, since the death forms of loser cells were no longer confined to apoptosis, but has presented different codes such as necroptosis, autophagy, entosis, cell-independent extrusion and cell senescence in cell competition (Bondar & Medzhitov, 2010; Hogan *et al*, 2009; Li & Baker, 2007; Ohsawa *et al*, 2011; Sun *et al*, 2014b; Villa Del Campo *et al*, 2014), the specific death form of lenvatinib-sensitive cells is required to be clarified in the future.

In recent decades, multi-omics analyses and numerous studies have indicated that mitophagy not only facilitates multiple malignance behaviors (Maes *et al*, 2014; Sun *et al*, 2014a; Wu *et al*, 2015) and tumor heterogeneity (Wang *et al*, 2022), but also induces resistance to anti-cancer drugs including doxolucibin, cisplatin, sorafenib and lenvatinib via a variety pathways and core molecules such as ARIHI, PINK1, NIX and BNIP3 (Villa *et al*, 2017; Wu *et al*., 2020; Yan *et al*, 2017; Zheng *et al*, 2021). Crucially, mitochondrial homeostasis and glycolysis activities including glucose uptake and lactic acid production also are regulated by mitophagy-related proteins (O’Sullivan *et al*., 2015; Springer *et al*., 2021; Zhang *et al*, 2011). However, the mechanism in mitophagy, chemoresistance and cell competition hitherto remains unknown. In this study, combined with further transcriptome analyses, experiment results of high-expressed BNIP3 and the colocalization or protein expressions of Cy5.5-LC3B and TOMM20 imply that BNIP3-mediated mitophagy induces energy metabolism switching and competitive fitness of lenvatinib-resistant cells in cell competition. Additionally, based on in-vitro *BNIP3* gene knockdown, reduced glycolysis activities and related proteins expression, especially key enzyme ENO2 and competitive advantage, which are rescued by transient overexpression of BNIP3. These results not only further support our conclusion, but also suggest the effect of mitophagy on glycolysis in lenveatinib-resistant cells is targeted by ENO2, which has been verified in various cancers (Kim *et al*., 2019; Sun *et al*., 2021). Although subsequent results under dual gene modification of ENO2 and BNIP3 further validate and reveal the terminal target of ENO2, we remain unclear why BNIP3-mediated mitophagy is specifically associated with ENO2 and importantly, how to connect with it in lenvatinib-resistant cells. As we know, under the limited source or stressful environment, metabolic alteration or shifting is bound to induce energy imbalance, and according to studies, the metabolic sensor AMPK subsequently is activated resulting from the rise of AMP/ATP ratio, eventually targeting glycolytic enzyme (Domenech *et al*, 2015). Therefore, through combining with clinical HCC cohorts, results of significantly enriched AMPK pathway, high-expressed phosphorylated AMPK and AMPK-IN-3 inhibitor-induced inhibition in glycolysis activities and competitive dominance of lenvetinib-resistant cells indeed prove that AMPK serving as signal sensor plays an essential role in encouraging glycolysis and maintaining winner status of lenvetinib-resistant cells in cell competition. Moreover, increased ENO2 expression and attenuated inhibitory effects of ENO2 knockdown on glycolysis activities under AMPK activator, unchanged BNIP3 expression and mitophagy colocalization under AMPK inhibiting or AMPK activating, decreased inhibitory influence of AMPK-IN-3 on AMPK expression and glycolysis activities and population advantage under BNIP3 overexpression and previous BNIP3-ENO2 regulatory axis first reveal a vital role for BNIP3-AMPK-ENO2 crosstalk in continuously maintaining HCC lenvatinib-resistant cells’ competitive dominance. Of note, it has been reported that AMPK modulates energy and metabolic homeostasis through phosphorylating key glycolytic enzyme PFKFB3 and on the other hand, AMPK-induced recycle signal can be effectively activated to devour cell debris and then promote the growth of lung cancer (Eichner *et al*, 2019); therefore, the detailed modification of AMPK-ENO2 axis and the effects of AMPK on lenvatinib-sensitive cells are required to be clarified in the future.

In summary, mitophagy can profoundly boost shifting energy production from mitochondrial oxidative phosphorylation to glycolysis via BNIP3-AMPK-ENO2 crosstalk, thereby maintaining lenvatinib-resistant cells’ growth and leading to HCC development through sacrificing lenvatinib-sensitive cells in cell competition. As cell competition driven by energy metabolism differences is notoriously involved in tumorigenesis and progression, and an increasing literature suggests a pivotal role for metabolic shifting-mitophagy in facilitating drug resistance and a variety of malignant behaviors (Sun *et al*., 2014a; Wang *et al*., 2022; Wu *et al*., 2015), we propose that cell competition serving as an essential tumor heterogeneity is expected to become a treatment basis and targeted mitophagy-related BNIP3 can be a promising strategy for overcoming chemoresistance.

## Materials and Methods

### Reagents and Tools Table

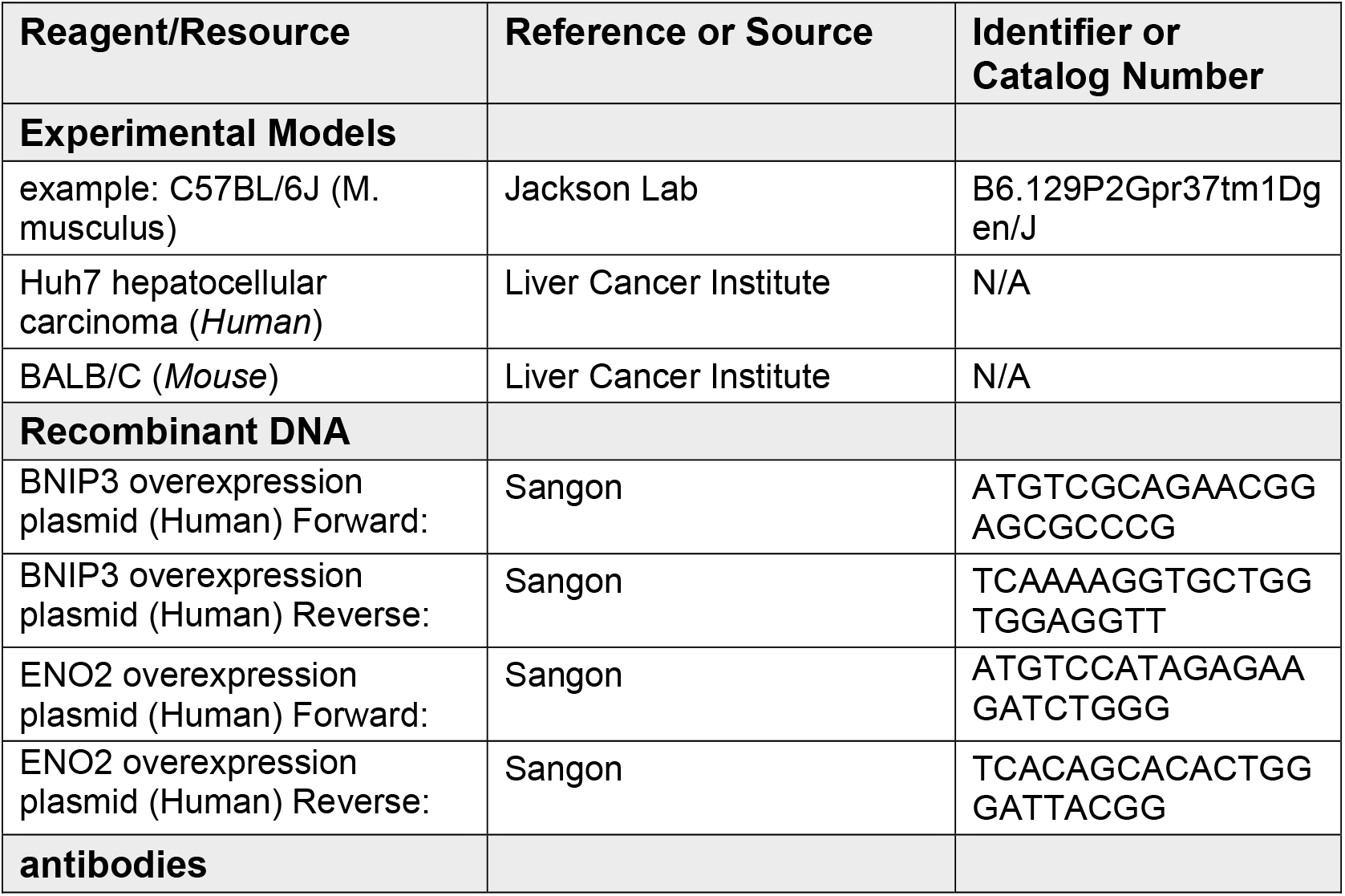
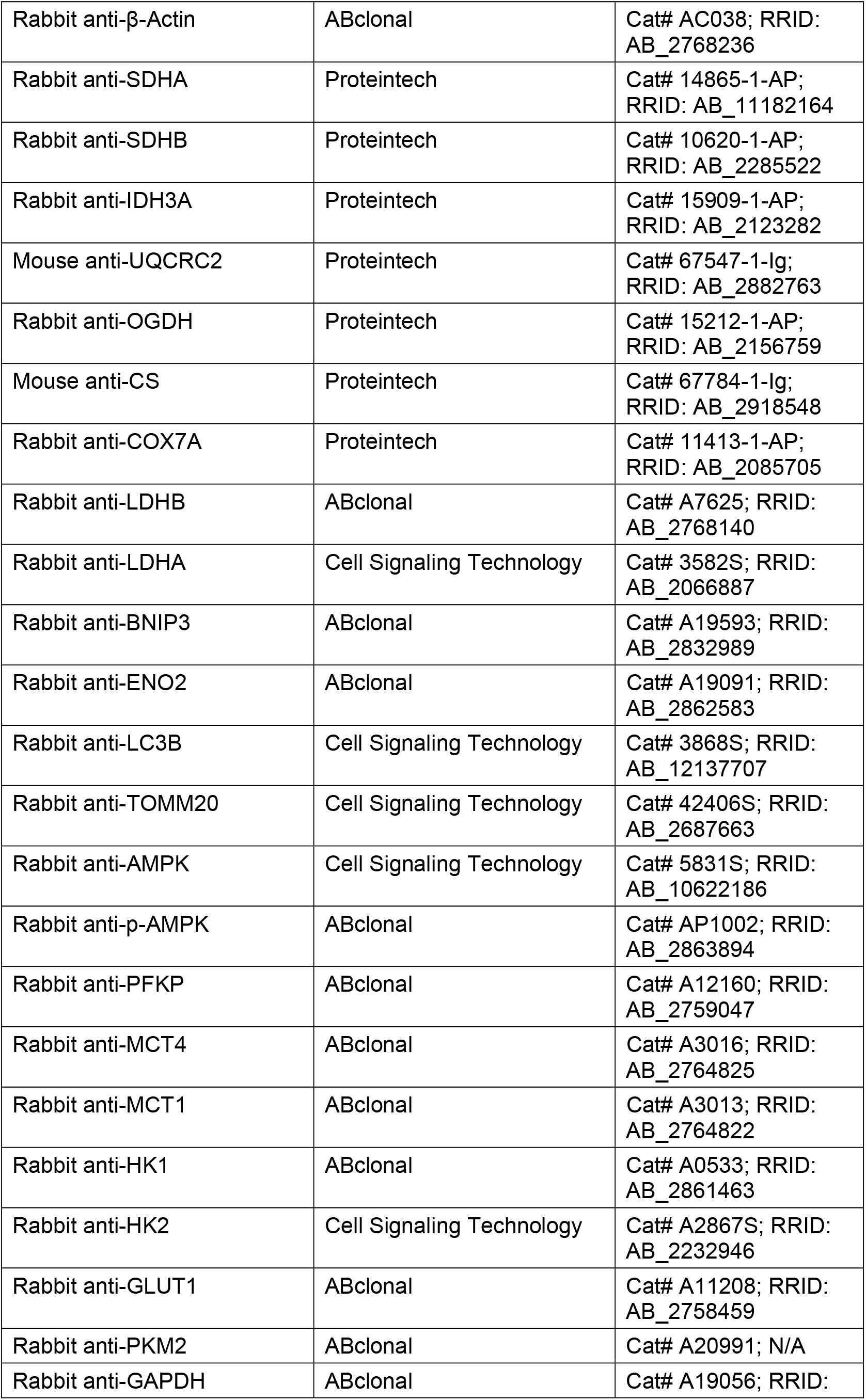
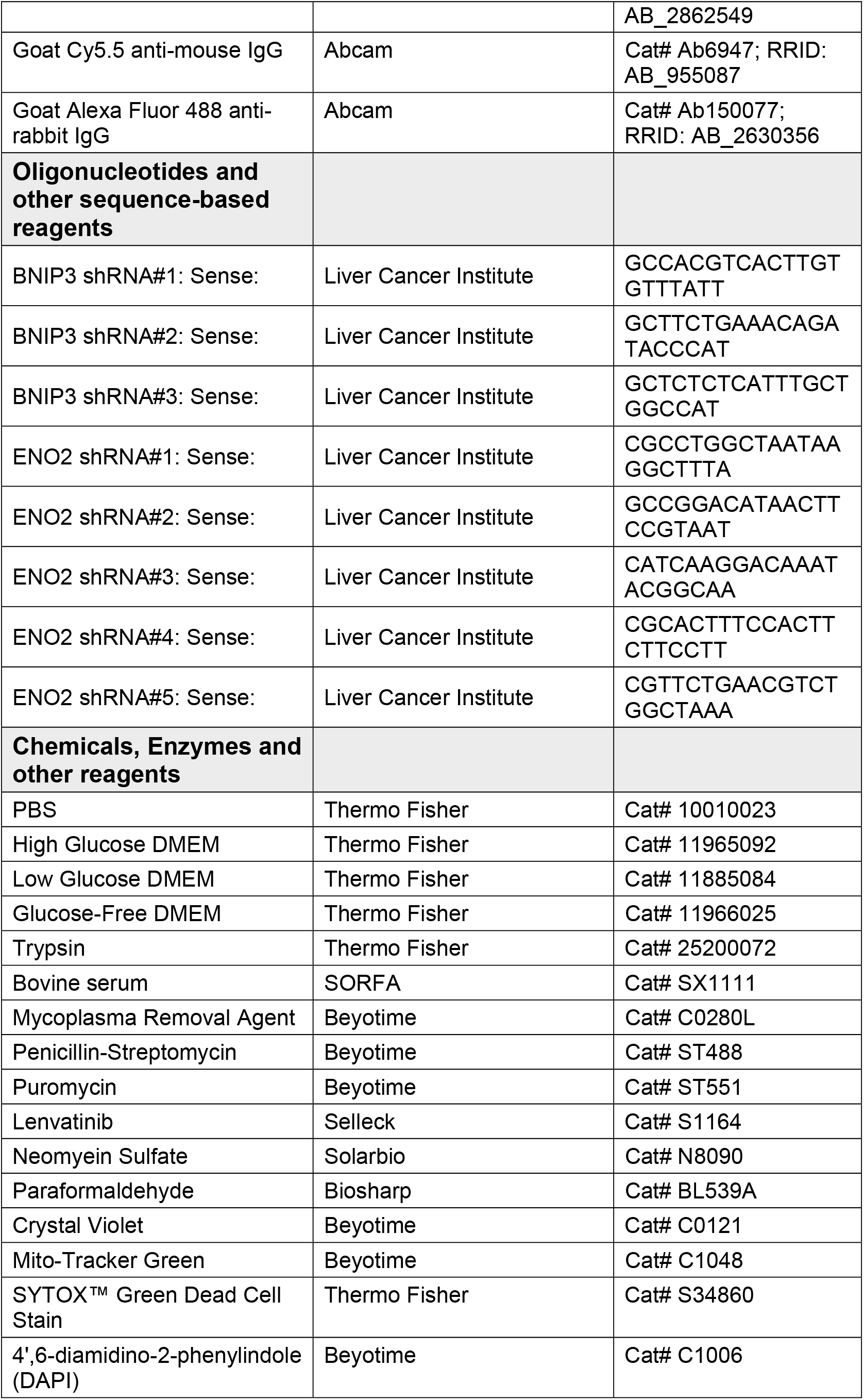
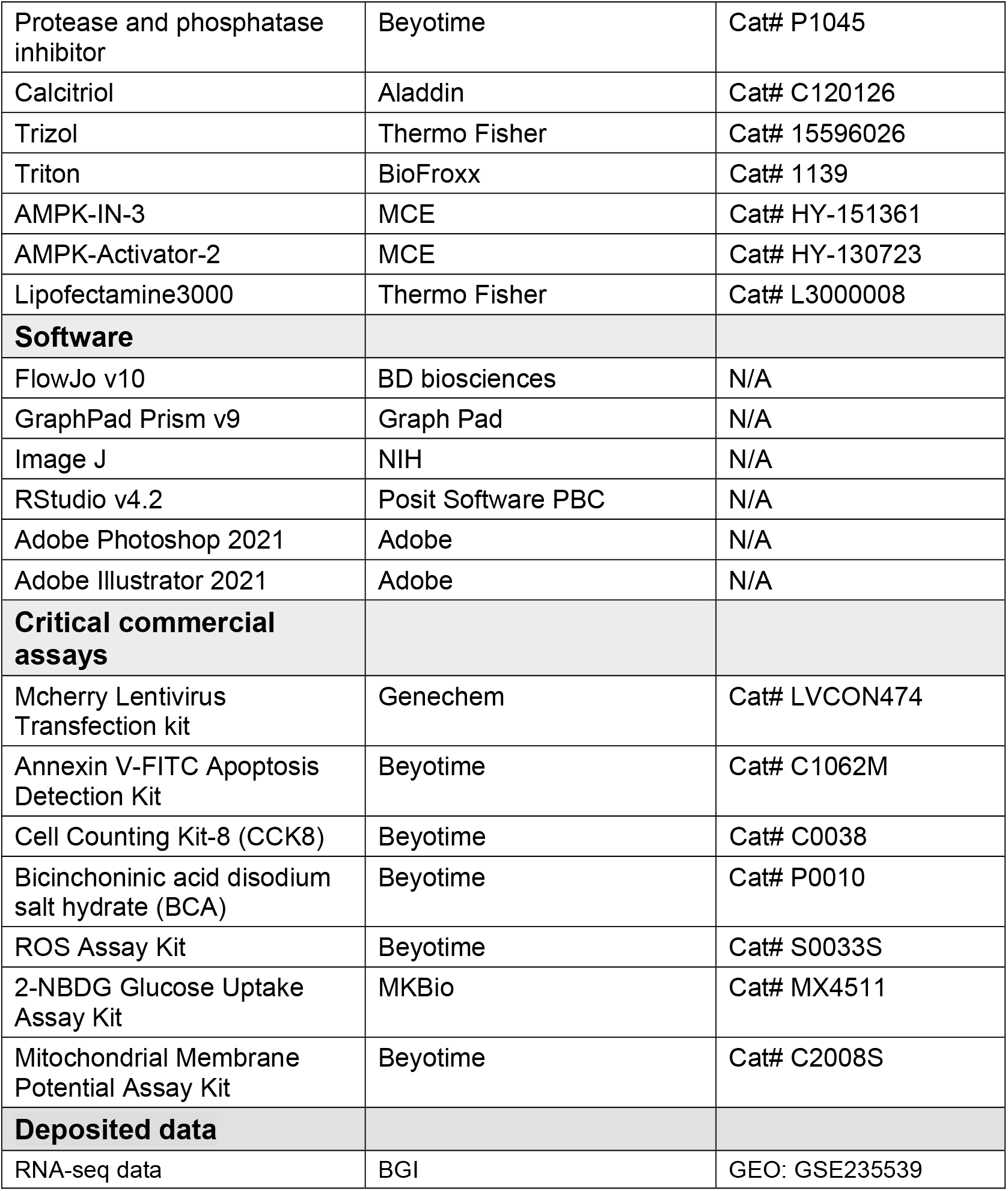

### Methods and Protocols

#### Cell lines

Human liver tumor cell lines (Huh7) obtained from the Liver Cancer Institute, Zhongshan Hospital, Fudan University (Shanghai, China) were cultured in High glucose DMEM (Thermo Fisher, USA) with 10% fetal bovine serum (SORFA, China), 1% penicillin and streptomycin (Beyotime China). The lenvatinib-resistant cells (Huh7R) were established from Huh7 cells by exposure to gradually increasing concentrations of Lenvatinib (MCE, USA). The final highest concentration and maintenance dose of Huh7R were 30μm and 20μm, respectively. To track or differentiate cells, Huh7 stably expressing mcherry (Huh7m) were constructed via transfecting Huh7 with CMV-MCS-EF1a-mCherry-T2A (Genechem, China) and following the selection of DMEM containing 5ug/ml puromycin (Beyotime, China). All cells were maintained at 37 °C with 5% CO2.

#### Mice

6-week-old BALB/c nude mice were housed and raised at laboratory animal center of Zhongshan Hospital, Fudan University (Shanghai, China), following the guidelines of biosafety and bioethics. To obtain tumor models based on coculture system, total 1X10^7^ cells of Huh7m mixed with Huh7/Huh7R at 1:1 ratio were injected subcutaneously to our BALB/c mice. All BALB/c mice were monitored and those tumor-carried BALB/c mice were assigned to our experimental groups. Subsequent fluorescence imaging and analyses were performed based on H&E-tissue staining of subcutaneous tumors which were obtained from experimental groups.

#### Cell Competition Assay

To visually observe cell competition behavior in vitro, Huh7/Huh7R were cocultured with Huh7m (NCC group, CC group) at 1:1 ratio, or Huh7R, Huh7 and Huh7m (Alone group) were cultured alone overnight. After cell attachment, time-lapse observation experiment was executed for 48h in 1ml medium per well by high content imaging system (Perkin Elmer Operetta). Images were exposed for 20ms in the mcherry channel and captured every thirty minutes for 48h using a 10X/20X objective lens. For flow cytometry cell competition analyses, Huh7/Huh7R were cocultured with Huh7m (NCC group, CC group) at 1:1 ratio, or Huh7R and Huh7m (Alone group) were cultured alone. After adherence, cells were collected at 24h/48h through trypsin (Thermo Fisher, USA) digestion and 1X PBS (Thermo Fisher, USA) resuspension. Specifically, Alone groups were manipulated as follows for comparison: collected Huh7R cells were mixed with collected Huh7m cells. All cells were measured by BD FACSCanto II (BD Biosciences). The experiments were carried out independently three times.

#### shRNA/plasmid transfection and chemicals treatment

To analyse the molecular mechanism of cell competition, Huh7R stably expressing *BNIP3*/*ENO2*-short hairpin RNA (sh*BNIP3*/*ENO2*) was constructed via transfecting cells with sh*BNIP3*/*ENO2* lentivirus obtained from the Liver Cancer Institute and following the selection of DMEM containing 5ug/ml puromycin; Huh7R transitorily overexpressing *BNIP3* (oe*BNIP3*) was constructed via transfecting Lipofectamine 3000 reagent (Thermo Fisher, USA) with *BNIP3* plasmid (Sangon, China) and following the selection of DMEM containing 5ug/ml neomyein sulfate (Solarbio, China); AMPK-activated/inhibited Huh7R was constructed by pre-treating with 2μm AMPK-activator-2 (MCE, USA)/15μm AMPK-IN-3 (MCE, USA). For rescue assays, Huh7R-sh*BNIP3* was transitorily transfected Lipofectamine 3000 reagent with *BNIP3* expression plasmid and following the selection of DMEM containing 5ug/ml neomyein sulfate; Huh7R-sh*ENO2* was transitorily transfected Lipofectamine 3000 reagent with *ENO2* expression plasmid (Sangon, China) and following the selection of DMEM containing 5ug/ml neomyein sulfate, or treated with AMPK-activator-2; Huh7R-oe*BNIP3* was treated with AMPK-IN-3; Huh7R were pre-treated with AMPK-activator-2 before adding AMPK-IN-3; then all the above Huh7R cells were cultured alone or cocultured with CCHuh7m for 48h before performing corresponding assays. To validate the metabolic difference of CCHuh7R and CCHuh7m, CC group was treated with low glucose addition (Thermo Fisher, USA)/ 12.5 μg/mL Calcitriol (Aladdin, China) and then performed corresponding assays. Mycoplasma removal agent (Beyotime, China) was regularly applied to all cell lines.

#### Cytotoxicity Experiment

To further observed the drug sensitivity between Huh7R and Huh7, Huh7/Huh7R (1X10^4^ cells/100ul) were plated into 96-well plates at 100ul/well and treated with lenvatinib at various concentrations (0, 0.01, 0.1, 0.5, 1, 10, 20, 40 and 100μm) for 48h after attachment. Subsequently, medium was removed, and cells were incubated in the fresh medium mixed with CCK8 working fluid (Beyotime, China) for 1h at 37 °C. Cell viability was measured through detecting the absorbance at wavelengths of 450nm using microplate reader (BioTek, USA). The experiments were carried out independently three times.

#### Cell proliferation assay

To compare the short-term proliferative capacity of Huh7 and Huh7m, Huh7 (5X10^3^) and Huh7m (5X10^3^) were respectively seeded into 96-well plates for 24h/48h. Next, the complete media was removed, and cells were incubated in the fresh medium mixed with CCK8 working fluid for 1h at 37 °C after washing by 1X PBS. Cell viability was measured through detecting the absorbance at wavelengths of 450nm using microplate reader. The experiments were carried out independently three times.

#### Colony Formation Assay

To compare the long-term proliferative capacity of Huh7 and Huh7m, Huh7 (1X10^4^) and Huh7m (1X10^4^) were separately seeded into 12-well plates for 15 days. In the meantime, media was changed every five days after 1X PBS washing. Lastly, cells were fixed in 4% paraformaldehyde (Biosharp, China) and stained with crystal violet staining solution (Beyotime, China). Afterwards, all the colonies were counted and quantitative analyses. The experiments were carried out independently three times.

#### ROS Measurement

To measure the intracellular ROS level between Huh7 and Huh7m, Huh7 (3X10^5^) and Huh7m (3X10^5^) were respectively seeded into 12-well plates overnight. After adherence, cells were followed by trypsinization after staining with 5µm 2, 7-dichlorofluorescein diacetate (DCFH-DA; Beyotime, China) for 30 min at 37 °C (protect from degradation of light). Subsequently, cells were resuspended by 1X PBS resuspension and collected. DCFH-DA intensity was detected by the FITC-A channel using BD FACSCanto II (BD Biosciences). The experiments were carried out independently three times.

#### Early Apoptosis Assay

To contrast the early apoptosis in Huh7 and Huh7m, Huh7 (3X10^5^) and Huh7m (3X10^5^) were seeded into 12-well plates overnight. After adherence, cells were followed by trypsinization after staining with Annexin V-FITC binding buffer and Annexin V-FITC (Beyotime, China) for 20min at 37 °C (protect from degradation of light). Subsequently, cells were resuspended by 1X PBS resuspension and collected. Annexin V intensity was detected by the FITC-A channel using BD FACSCanto II (BD Biosciences). The experiments were carried out independently three times.

#### 2-NBDG Glucose Uptake Assay

Cells mentioned previously were cultured for 48h and stained with 100ul glucose-free medium (Thermo Fisher, USA) containing 200μm 2-NBDG (MKBio, China) and incubating at 37°C for 80min (protect from degradation of light). For flow cytometric analysis (BD Biosciences), cells were digested and resuspended by 1X PBS resuspension. 2-NBDG intensity was detected by the FITC-A channel. The above experiments were carried out independently three times.

#### Mitochondrial Membrane Potential Assay

Cells mentioned previously were cultured for 48h and stained with 0.1X Rhodamine123 (Beyotime, China) and incubating at 37°C for 30min (protect from degradation of light). For flow cytometric analysis (BD Biosciences), cells were digested and resuspended by 1X PBS resuspension. Rhodamine123 intensity was detected by the FITC-A channel. The above experiments were carried out independently three times.

#### Dead Cell Detection Assay

Cells mentioned previously were cultured for 48h and followed by trypsinization after staining with SYTOX Green (Thermo Fisher, USA) and incubating at 37°C for 30min (protect from degradation of light). Subsequently, cells were resuspended by 1X PBS resuspension and collected. SYTOX Green intensity was detected by the FITC-A channel using BD FACSCanto II (BD Biosciences). The experiments were carried out independently three times.

#### Mitochondrial morphology Assay

Cells mentioned previously were cultured for 48h and stained with 1:1000 Mito-Tracker Green (Beyotime, China) and incubating at 37°C for 90min (protect from degradation of light). Two cellular samples were prepared for different experiments. For high content fluorescence imaging (Perkin Elmer Operetta), cells were also stained with and 1X DAPI for 10min and added in 1X PBS after removing the mixture and analyzed by the Harmony software (Perkin Elmer) using a 63X water immersion objective lens. For flow cytometric analysis (BD Biosciences), cells were digested and resuspended by 1X PBS resuspension. The above experiments were carried out independently three times.

#### Western blot

Cells mentioned previously were cultured for 48h; then cells of coculture groups were divided by flow sorting (BD Biosciences) based on mcherry fluorescence expression. Afterwards, cells of all groups were separately lysed in 1X RIPA cell lysis buffer containing 1X protease and phosphatase inhibitors (Beyotime, China). BCA assay was then performed to determine collected protein concentrations. A SDS-PAGE loading buffer was added to our samples and heated at 100°C for 5min. In the next step, samples were executed for 10% SDS PAGE and western transfer. Subsequent 1h block by western sealing buffer and primary antibodies were performed at 4°C overnight. Primary antibodies utilized in this paper were including β-Actin (ABclonal, USA), ENO2 (ABclonal, USA), GLUT1 (ABclonal, USA), HK1 (ABclonal, USA), HK2 (Cell signaling technology, USA), MCT1 (ABclonal, USA), MCT4 (ABclonal, USA), PFKP (ABclonal, USA), PKM2 (ABclonal, USA), LDHA (Cell signaling technology, USA), LDHB (ABclonal, USA), GAPDH (ABclonal, USA), LC3B (Cell signaling technology, USA), TOMM20 (Cell signaling technology, USA), BNIP3 (ABclonal, USA), AMPK (Cell signaling technology, USA), p-AMPK (ABclonal, USA), IDH3A (Proteintech, USA), COX7A (Proteintech, USA), SDHB (Proteintech, USA), SDHA (Proteintech, USA), UQCRC2 (Proteintech, USA), OGDH (Proteintech, USA) and CS (Proteintech, USA). After washing 3 times by PBST buffer, membranes were incubated at 37°C for 1h by secondary antibodies. At last, chemiluminescence was used for detecting protein levels of membranes. The above experiments were carried out independently three times.

#### Mitophagy Colocalization Assay

Cells mentioned previously were cultured for 48h and fixed in 4% paraformaldehyde and then permeabilized in 0.5% Triton X-100 for 15min to improve antigen accessibility. After permeabilization, 30min block by 5% BSA and primary antibodies specific for LC3B, TOMM20 were performed at 4°C overnight. After washing 3 times by PBST buffer, cells were incubated at 37°C for 1h by secondary antibodies including Cy5.5 goat anti-mouse IgG (Abcam, UK) and Alexa fluor 488 goat anti-rabbit IgG (Abcam, UK) (protect from degradation of light). At last, cells were analyzed by the Harmony software (Perkin Elmer) using a 63X water immersion objective lens after washing by PBST buffer. The above experiments were carried out independently three times.

#### RNA isolation and Libraray preparation

Huh7R/Huh7 and Huh7m were cocultured in a 1:1 ratio for 48 hours. Flow sorting (BD Biosciences) was used to separate the different cell populations in the coculture sample based on mcherry expression. Three biological replicates for each of the different culture condition of Huh7, Huh7m and Huh7R were performed. Equivalent cells were collected, and total RNA of the cells were extracted by TRIzol reagent (Thermo Fisher, USA) conducting the same method in RT-qPCR experiment. A total of 2 µg RNA per specimen was ready for RNA sequencing experiment. Agilent 2100 Bioanalyzer (ABI, USA) was performed to assessed quantity and quality of the cDNA libraries. The Double-stranded PCR products were treated with thermal denaturation and cyclization reaction using the splint oligo sequence and the final library was generated. Phi29 (Thermo Fisher, USA) was utilized for amplifying the single-stranded circular DNA and then a DNA nanoball was generated.

#### RNA-seq and differential expression analysis

MGISEQ-2000 platform was used to sequence the prepared libraries mentioned above. SOAPnuke (Li *et al*, 2008) (Version 1.5.2) was performed to clean the sequencing data via removing reads which contained sequencing adapter, contained more than 20% of low-quality base ratio (ratio less than or equal to 5) and contained more than 5% unknow base (‘N’ base) ratio. The cleaned reads, consequently, were acquired and stored in FASTQ format. Then, HISAT2 (Kim *et al*, 2015) was used to align the cleaned reads to the reference human genome. Bowtie2 (Langmead & Salzberg, 2012) (Version 2.2.5) was used to align the cleaned reads to the reference coding gene set. Afterwards RSEM (Li & Dewey, 2011) (Version 1.2.12) was used to compute the expression level of gene and Count data was generated. Essentially, differential expression analysis (DEA) for Count data was executed using the Limma-Voom algorithm of R-package Limma (Version 3.52.1). Selection criteria of significant differentially expressed genes (DEGs) was as follows: p-value < 0.05, FDR < 0.05 and |Log2FC| > 1. Transcripts per million (TPM) normalization was used to scaled expression per transcriptome.

#### Data resource and analysis

RNA-seq data of 374 HCC samples were obtained from The Cancer Genome Atlas-Liver Hepatocellular Carcinoma (TCGA-LIHC) cohort downloading from the UCSC Xena website (http://xena.ucsc.edu/). RNA-seq data of 240 HCC samples were acquired from International Cancer Genome Consortium-Liver Hepatocellular Carcinoma (ICGC-LIHC) cohort. The annotation information data (gencode.v22.annotation.gene.probeMap) downloaded from UCSC Xena website was performed to match the Ensemble Gene IDs to Gene Symbols. According to the median value of BNIP3 mRNA expression, TCGA-LIHC cohort and ICGC-LIHC cohort were respectively divided into BNIP3 high-expression group and BNIP3 low-expression group. Afterwards differential expression analysis, Pearson correlation analysis and Gene set enrichment analysis were executed in TCGA-LIHC cohort and ICGC-LIHC cohort, respectively.

#### Gene set enrichment analysis (GSEA)

To explore biological characteristics in indicated comparative groups respectively, GSEA was performed to score the gene sets based on the differential expression analysis via using R-package clusterProfiler (Version 4.4.2). Genes rank in the indicated gene sets was utilized in accordance with results of differential expression analysis within CCHuh7R vs Huh7R group, CCHuh7R vs NCCHuh7 group, Huh7R vs Huh7 group and BNIP3 high-expression vs BNIP3 low-expression group in two cohorts mentioned above. Customized gene set composed by 38 glycolysis-related pathways based on KEGG database (http://www.genome.jp/kegg/) was performed to explore the regulatory mechanism between Mitophagy and Glycolysis in our indicated groups. Selection criteria of significant biological characteristics was as follows: p-value < 0.05, FDR < 0.05 and |NES| > 1.

#### Gene set variation analysis (GSVA)

To investigate the mitochondria mass and activities in indicated comparative groups respectively, GSVA was carried out on log2 (normalized TPM +1) expression values within CCHuh7R vs Huh7R group and CCHuh7R vs NCCHuh7 group via using R-package GSVA (Version 1.44.1). Customized gene set composed by pathways of mitochondria mass and activities based on GO database (http://geneontology.org/page/go-database) was performed to explore indicated phenotypes. GSVA enrichment score was generated for each sample and pathway in indicated groups. Afterwards, DEA was executed and, selection criteria of significant differentially activities were as follows: p-value < 0.05, FDR < 0.05.

#### Single sample GSEA (ssGSEA)

To examine the association of BNIP3 with certain pathways, ssGSEA was carried out on log2 (normalized TPM +1) expression values within BNIP3 high expression vs BNIP3 low expression group in two cohorts mentioned above via using R-package GSVA (Version 1.44.1). The gene sets for AMPK signaling pathway and Glycolysis pathways were obtained in GSEA results based on customized gene set mentioned above. Enrichment score was generated for each sample in two cohorts, respectively. Afterwards, Pearson correlation analysis was implemented to determine correlation between BNIP3 expression and AMPK expression or two pathways described above via using R-package ggpmisc (Version 0.4.6).

#### Data visualization statistics

Volcano plots were generated based on DEGs in indicated comparative groups. Heat maps were generated for indicated pathways-related DEGs or pathways based on results of GSEA and GSVA. UpSet plots were generated for Top 50 genes or up-regulated genes of glycolysis pathway based on DEGs and GSEA in indicated comparative groups via using R-package UpSetR (Version 1.4.0). Pearson’s correlation coefficient was conducted to determine the correlation between BNIP3 expression and AMPK expression or two pathways described above in indicated groups. Visualization of analyses mentioned previously was plot by using ggplot2 R package. All of analyses and visualizations were performed in the R software (Version 4.1.3) using multiple R packages. All the statistical analyses were performed via carrying out GraphPad Prism 9.0 software.

## Data availability

The access number of the processed expression datasets of RNA-seq from this paper is GSE235539.

No original code was reported in this paper.

Any other data in this paper are available upon reasonable request for the corresponding author.

## Acknowledgments

We thank all collaborators at Zhongshan Hospital of Fudan University for helping and supporting this study. This study was funded by the National Natural Science Foundation of China (81872492).

## Author contributions

**Sikai Wang:** Conceptualization; experimental design; investigation; formal analysis; validation; sequencing data analysis; visualization; initial draft; editing and finalization. **Hongxia Cheng:** Conceptualization; experimental design; investigation; formal analysis; validation; visualization; initial draft. **Miaomiao Li:** Investigation. **Haoran Wu:** Validation. **ShanShan Zhang:** Validation. **Dongmei Gao:** Validation. **Yilan Huang**: Validation. **Kun Guo:** Conceptualization; experimental design; visualization; editing and finalization; funding support; resources; supervision.

